# A Bayesian Multi-Species Approach Infers Gene Regulatory Networks Across Non-Model Organisms

**DOI:** 10.64898/2026.08.25.746862

**Authors:** Andrew Soborowski, Omur Kayikci, Mar Martinez Pastor, Julie A. Maupin-Furlow, William H. Majoros, Amy Schmid

**Author notes:** These authors also contributed equally to this work. Deceased. Membership list can be found in the Acknowledgments section.

## Abstract

Control of gene expression by transcription factors (TFs) is a critical mechanism for cells to maintain homeostasis in response to environmental signals. Gene network models that predict regulatory interactions between transcription factors and the genes they control aid in understanding these complex processes. These models are useful as they provide testable hypotheses of regulatory interactions, transcription factor function, and accelerate the study of uncharacterized transcription factors. However, inference of these models is computationally challenging due to the vast quantity of data required given the many possible states of the regulatory network. Microbial genomes encode hundreds of transcription factors, with numerous interactions that require substantial functional genomics datasets to infer. This problem is accentuated in understudied organisms, species that would greatly benefit from an inferred network for biological discovery, where the lack of available data is particularly constraining for effective inference. To address this problem, we have developed GRN-BMuSeR (Gene Regulatory Networks from Bayesian MUlti-SpEcies Regression), a novel multitask approach to gene regulatory network inference that leverages gene orthology between closely related species to improve inference performance. We evaluate its performance on a dataset from the well-studied bacterial species *Bacillus subtilis*, demonstrating improved performance in multitask settings. Applying the model to simulated data reveals utility in multi-species contexts. Finally, we apply our models to infer GRNs and explore predictions for two hypersaline-adapted archaeal species. We leverage a rich dataset from *Halobacterium salinarum* to inform the inference of the gene regulatory network of *Haloferax volcanii*, for which a more limited genomics dataset was available. We generate a large compendium of gene expression data for *Hfx.volcanii* for GRN inference input. Through exploration of resultant network predictions, we show concordance with known TF functions and discover hundreds of novel TF functional predictions. Moving forward, our results provide a framework to generate testable hypotheses that will serve to guide experimental work and accelerate discovery in these understudied species.

## Introduction

Cellular survival requires a timely and robust response to environmental stress. An organism’s capacity to respond to stress defines its niche as well as its resilience to both transient and permanent environmental shifts. Additionally, study of microbial stress responses has led to an elucidation of protein functions and increased knowledge of cellular physiological processes [1, 2]. In response to environmental cues, transcription factors (TF) serve to regulate gene expression by promoting or inhibiting the binding of RNA polymerase, regulating the production of mRNA for a given gene.

Inferring a gene regulatory network (GRN) to conceptualize and predict the interactions between TFs and the genes they regulate facilitates an understanding of how organisms adapt to shifting environmental conditions [3–6]. GRN models represent TFs and genes as nodes in a graph, with edges between TFs and genes representing a regulatory interaction. GRN inference involves ranking these TF-gene interactions using gene expression data, with potential incorporation of orthogonal datasets such as TF-DNA binding activity data (ChIP-seq) to boost performance [7]. GRNs have been used previously in model microorganisms to study gene regulation on a global scale and understand cell fates and regulatory paradigms in multicellular organisms [8]. In understudied species such as the microorganisms of the domain Archaea, inferred networks have been used to suggest downstream perturbation experiments and to generate hypotheses for TFs with no or little known function [9, 10]. For archaea, GRNs are a considerable aid to experimental design when a large fraction of the genome function remains unexplored. Understanding the GRNs of these organisms is of keen interest due to their important contributions to biotechnology and global geochemical cycles [11, 12]. Although GRN models have been successful in generating hypotheses for rapid discovery of TF environmental response functions in archaea [13, 14], they were inferred for single species, leading to questions of whether these regulatory relationships are conserved. To facilitate inference in understudied but environmentally important microorganisms, here we design and implement a multi-species GRN learning framework using multitask machine learning.

Multitask inference is a machine learning approach where a model learns from two or more related tasks, sharing common information to allow for a better fit in each individual task [15]. This framework is not new to GRN inference, where it has been successfully applied to correct for batch effects across disparate datasets and for the joint inference of networks across cell types and developmental stages [16]. However, previous multitask inference methods are limited to a single species because they require one-to-one identity of genes in the underlying regulatory networks across tasks [16].

While portions of the network may differ due to context specific expression captured only in a subset of tasks, the high degree of similarity between the remainder of the network allows for improved inference in a multitask setting. Other models either infer individual networks separately and compare TF-gene connections post-hoc [17], exclude non-orthologous genes [18], or require additional orthogonal data and assumptions that may not be available for all species [19]. Therefore, a fundamental challenge remains to jointly infer GRNs across organisms for which only a subset of genes are homologous.

Here, we address this challenge by expanding upon the multitask framework introduced in previous approaches that required identical nodes across tasks [16]. We relax the assumption that each task must be derived from observations from the same species. We propose the GRN-BMuSeR (Gene Regulatory Networks from Bayesian MUlti-SpEcies Regression) model that jointly constructs networks between species, leveraging the available data in one organism to drive the inference of a GRN model in a related, understudied species. Our approach relies upon and extends the regularized horseshoe prior into the multitask setting. Information flows between tasks in the context of the horseshoe prior’s imposed sparsity [20] in the case of homologous genes and TFs, while allowing inference to proceed in the case of unique genes and TFs. As a practical application of GRN-BMuSeR, we infer two genetically tractable hypersaline-adapted archaeal species *Haloferax volcanii* and *Halobacterium salinarum*. Based on 16S rRNA sequence identity, these organisms are estimated to have diverged approximately 595 million years ago [21–23]. Phylogenetically, they are classified within the same Order but in different Families [24].

Archaea dominate in extreme environmental conditions that are often anathema to bacteria and eukaryotes, including extreme temperature, pH, and salinity environments [25–28]. Thriving in extreme salinity environments, haloarchaea (halophiles) serve as tractable model organisms for understanding archaeal genetics, genomics, physiology, and stress response due to their relative ease of growth and wealth of resources for genetic modification across multiple related species [29–33].

Historically, the bulk of GRN research efforts in halophiles (and archaeal organisms more generally) have focused on the extremely halophilic organism *Halobacterium salinarum* [9, 14, 34–36]. In the last decades, focus has shifted to studying *Haloferax volcanii*, a more moderate halophile, due to its rapid growth, facile genetic tools, and strong research community [37]. Both organisms are “salt-in” strategists that accumulate molar levels of intracellular potassium to combat osmotic imbalance in hypersaline lakes and salt evaporation ponds that approach sodium chloride saturation (~30%). As a result, the proteomes of *Hfx. volcanii* and *Hbt. salinarum*, like those of other extreme halophiles, are enriched in acidic residues [38]. However, in addition to constant salt stress, the lakes these organisms dwell in are often subject to harsh and rapid swings in temperature, radiation, and nutrient stress, each of which requires rapid and robust inducible stress responses [39]. Thus, halophiles present an opportunity to understand how organisms already adapted to extreme conditions adjust their physiology during environmental fluctuations.

In this work, we demonstrate that GRN-BMuSeR is able to perform comparably to existing multitask approaches on the same-species problem. Through simulations, we demonstrate that our approach remains effective when the same-species assumption is relaxed and the tasks are allowed to diverge from each other. We then apply our method to infer joint networks between halophilic archaeal species. We generated 149 RNA-seq profiles for 34 *Hfx. volcanii* TF mutants across 5 conditions to drive network inference. Using this dataset and leveraging the large existing transcriptomics compendium for *Hbt salinarum*, we demonstrate the utility of the method for effective inference of GRNs for data-poor species [40]. Using gene functional enrichment of network neighborhoods, we show that the network recapitulates known TF functions as well as enables prediction of potentially novel TF functions.

## Materials and methods

### 0.1 Generating RNA-Seq Data for *Hfx. volcanii*

Strains listed in Supplementary Table S1 were streaked from glycerol freezer stocks onto YPC18 [41] agar plates and incubated for 5 days at 42°C. Three to four single colonies of each strain were inoculated into 3 ml YPC18 liquid medium and grown to stationary phase overnight to synchronize the growth (optical density at 600 nm [OD600] ~ 1.0-2.0). Cells were then washed three times with basal salt solution [42] and transferred into 25 ml fresh YPC18 or HvCA liquid medium [41] with or without glucose (0.1%) with an initial OD600 of 0.05, and grown aerobically with 225 RPM rotary shaking at 42°C. Cells were harvested at mid-exponential phase (OD600 0.3-0.8) and total RNA was extracted using Absolutely RNA Miniprep kits (Agilent Technologies; Cat. No. 400800) or Quick-RNA Miniprep (Zymo Research; Cat. No. R1054). RNA was quantified and shown to be free of contaminating genomic DNA using the Agilent Bioanalyzer RNA Nano 6000 chip (Agilent Technologies, Cat. No 5067-1511). Ribosomal RNA was depleted from total RNA using the *Haloferax volcanii* riboPOOL kit (siTOOLs Biotech, Cat. No. dp-K024-016) according to reference [43]. RNA sequencing libraries were generated using 10-20 ng mRNA as input with the KAPA RNA Hyper Prep Kit (Cat. No. KK8541/ Roche: 08098107702) or the Watchmaker mRNA Library Prep Kit (Watchmaker Genomics, Cat. No.7BK0001) following the manufacturer’s directions. The fragment size of the libraries was measured using the Agilent 5300 Fragment analyzer and then pooled and sequenced on Illumina NovaSeq X Plus at the Duke Sequencing and Genomic Technologies Facility at Duke University (Durham, NC).

### 0.2 Analyzing RNA-seq Data and Gathering Species Compendia

Raw reads for the compendium using the previous experimental approach and gathered from previously published experiments housed in the National Center for Biotechnology Information (NCBI) Gene Expression Omnibus (GEO) database (Supplementary Table 1) [42–50]. Raw reads were trimmed and quality filtered using fastp [51] under default parameters with results visualized using multiQC [52]. Reads were aligned to the *Hfx. volcanii* DS2 reference genome (RefSeq Version 2.7.4A) using STAR [53] and counts tabulated using HTSeq [54]. Counts were normalized using the DESeq2 median of ratios method [55]. For *Hbt. salinarum*, a previously described compendium of normalized transcriptomics data comprised of 1,154 transcriptomics profiles across the 2,400 genes was used [40].

### 0.3 *Hfx. volcanii* RNA-seq Data Exploratory Analysis

The resultant dataset was log2 transformed to calculate Spearman correlation across samples using R. Heatmaps were generated using the pheatmap package in R with hierarchical clustering by Euclidean distance [56]. Fractional variance explained by each variable was calculated using the variancePartition package in R with each variable considered as a random effect in the context of a linear mixed model [57]. Under this model, random effects are estimated by maximum likelihood.

### 0.4 Hfx. volcanii and Hbt. salinarum TF Identification

TF coding genes were identified in the genomes of *Hfx. volcanii* and *Hbt. salinarum* and used in inference if they met the following criteria: (a) functional annotation in “transcription” from archaeal Clusters of Orthologous Genes (arCOG) ontology [58] and the NCBI annotation database (*Hfx. volcanii* RefSeq ID GCF 000025685.1 ASM2568v1); *Hbt. salinarum* RefSeq ID GCF 000006805.1 ASM680v1); (b) non-redundant encoding in the genome; (c) not associated with the core RNA polymerase machinery (i.e. these would be expected to regulate every gene, thereby violating sparsity assumptions, so were excluded); (d) functional prediction of membership in a TF protein family; (e) TFs not captured by these criteria but show orthology to an experimentally characterized TF the other species. This resulted in 158 TFs included in the inference for *Hfx. volcanii*, 87 for *Hbt. salinarum*. For *Hbt. salinarum*, the previously annotated list of 135 TFs [9] was included as a starting point, which was updated in the current study according to the aforementioned criteria. TFs of known function that were experimentally characterized according to the literature were added to the list if not captured by the above criteria.

### 0.5 Simulated Data Construction

We simulated pairs of differentially divergent networks from an ancestral state (for details, see GitHub code and Supp file 1). In brief, we first simulated an underlying ancestral regulatory network. We forced this network to diverge from itself into two descendants through the process of node (TF or gene) loss and gain as well as edge (TF-gene interaction) rewiring. The proportion of changes allowed between the resultant networks was controlled by a hyperparameter we called “divergence”. From the resulting descendant networks, we further simulated condition-based expression data under a noisy linear model based on the ground truth network structure. This expression data was used as input for GRN-BMuSeR to evaluate its ability to recapitulate the ground truth network as the descendant networks became increasingly diverged from each other.

### 0.6 GRN-BMuSeR

We modeled the expression of a given gene as a weighted sum of transcription factor expression values. The goal of this approach is to learn the weight terms under a sparse framework, where we interpret non-zero weights as predictions for gene-TF regulation. We implemented our approach as three distinct models: the default multitask model GRN-BMuSeR, a reduced model that runs on a single task GRN-BMuSeR ST, and an extended multitask model that incorporates task relatedness GRN-BMuSeR Ext.

#### 0.6.1 Primary Model

We modeled the expression *Y_i,j_*of gene *i* in condition *j* as the weighted sum of transcription factor *k* expression values (Eq. 1). Expression values for RNA-seq data were obtained as described above, while expression values for microarray data were obtained from their respective publications [16, 40]. In both cases, expression values were mean-variance normalized. In the case that sufficient data is available to infer transcription factor activity (TFA) values from prior data, we substituted *X_j,k_* with *A_j,k_*, where *A_j,k_* represents TFA as defined in [16].

#### 0.6.2 Definition of Tasks, Transcription Factors, and Homology

We defined task *t* as a distinct species and gene expression dataset to be included in the model. Each task was defined separately with its own list of target genes, list of TFs, and gene expression dataset. We specified that each gene may be modeled as a target, a TF, or both. Genes and TFs may be linked between tasks by identifying them as orthologs. For same-species inference, we defined orthologs as the same gene in each dataset. For multi-species inference, we defined a pair of genes as orthologs if they were the top hits of each other in a reciprocal BLAST search [59]. This ortholog mapping assumes that proteins sharing sequence identity are most likely to share function. Our model structure does not allow for one-to-many or many-to-many orthology relationships, so at most one ortholog across species was included in the model input for each gene or TF.

#### 0.6.3 Single task Bayesian Model

The core single task Bayesian model, implemented in GRN-BMuSeR ST, is applied in the case that only a single task is supplied or in the case that a given gene is unique to a certain task. In the single task setting, the core regression statement is given according to the following equation:

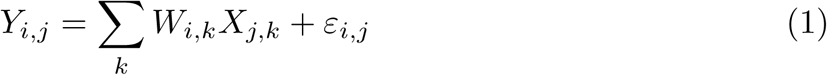

Here, *W* refers to a weight matrix of interactions between gene expression *Y* and TF expression *X*. This formulation implies a normal prior on *Y* given *W* and *X* with an implicit residual term given by *ε*. The full set of priors and definitions is given as follows where *ν*, *s*, and *P*_0_ are hyperparameters, *K* is total number of TFs and *J* is the total number of experimental conditions:

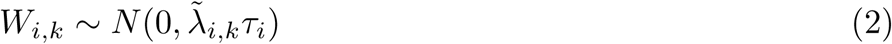

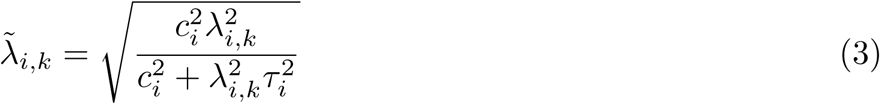

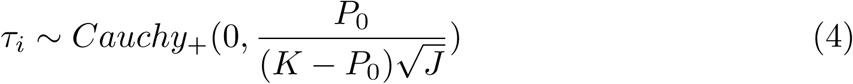

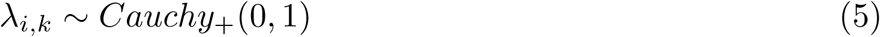

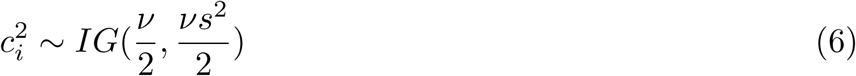

Above, eqs. (2-6) describes a regularized horseshoe prior placed on *W*. A detailed description of this approach is given in [20] but in brief, the prior shrinks the variance of each entry in *W*, and thus the sampled value of *W* (Eq. 2). The amount of shrinkage is calculated from both a global (*τ*) and local (*λ̃_i,k_*) level. This shrinkage restricts most values of *W* to near zero, in line with the assumption that most predictors are uninformative for a given gene. Meanwhile, strongly explanatory values *W* escape shrinkage to take on values distant from 0. The transformation in 3 truncates the unbounded Cauchy distribution defined in 5, greatly improving sampling efficiency and stability [20].

#### 0.6.4 Multitask Bayesian Model

The multitask model, implemented as GRN-BMuSeR, directly extends the single task model, incorporating the addition of multiple tasks while allowing for an incomplete ortholog linkage between genes and TFs across tasks. In this setup, the multitask regression model expands to the following:

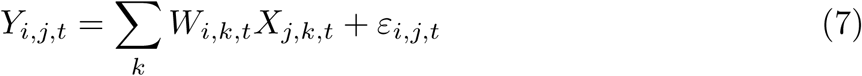

In this formulation, *i* indexes genes, *j* indexes conditions, *k* indexes TFs, and we introduce *t* which indexes tasks. Like before, the setup implies a normal prior on *Y* with an implicit residual term. The full set of priors for the multitask model expands to the following:

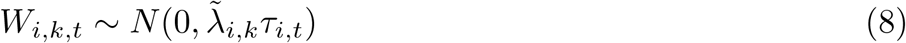

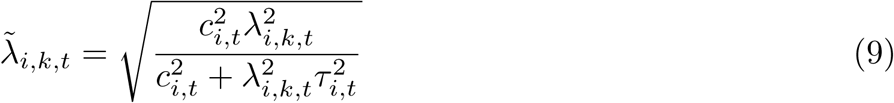

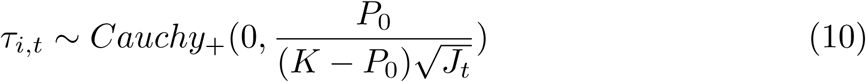

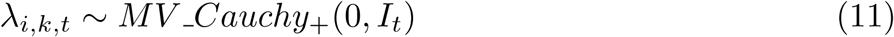

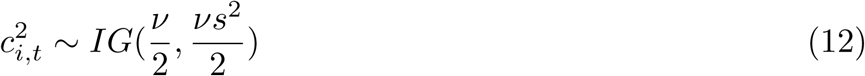

In the multitask model, separate regularized horseshoe priors are placed on the weight matrix for each task. For TFs that are homologous between tasks, *λ* values for each group of homologs is drawn from a multivariate Cauchy distribution with mean 0 and scale given by the identity matrix. While the draws from this distributed are uncorrelated, they not statistically independent, allowing evidence for a non-zero *λ* value in one task to influence the value of *λ* in other tasks. For TFs that do not have homologs in other tasks, values for *λ* are sampled from a standard Cauchy distribution as defined in 5.

#### 0.6.5 Multitask Bayesian Model Extension

To explicitly model task similarity, we propose the following extension to our method, implemented as GRN-BMuSeR Ext. In this extended multitask model, the prior distribution placed on the weights were instead altered by jointly sampling the weights of homologous TFs from a multivariate normal distribution given as:

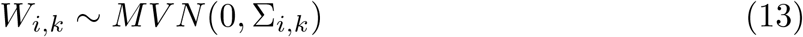

Here, the covariance matrix Σ associated with a given gene *i* and orthologous TFs *k* was decomposed into a vector of variances, *σ_i,k_*, and correlation matrix *ρ_i_* that describes correlation between task similarity at the level of each gene *i*. Variance values were calculated as described in the multitask model according to 8 while the correlation was inferred using an uninformative *LKJ* prior with hyperparameter *η*:

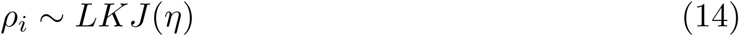

Graphical depiction of all three models are given in Fig 1.

**Fig 1.**
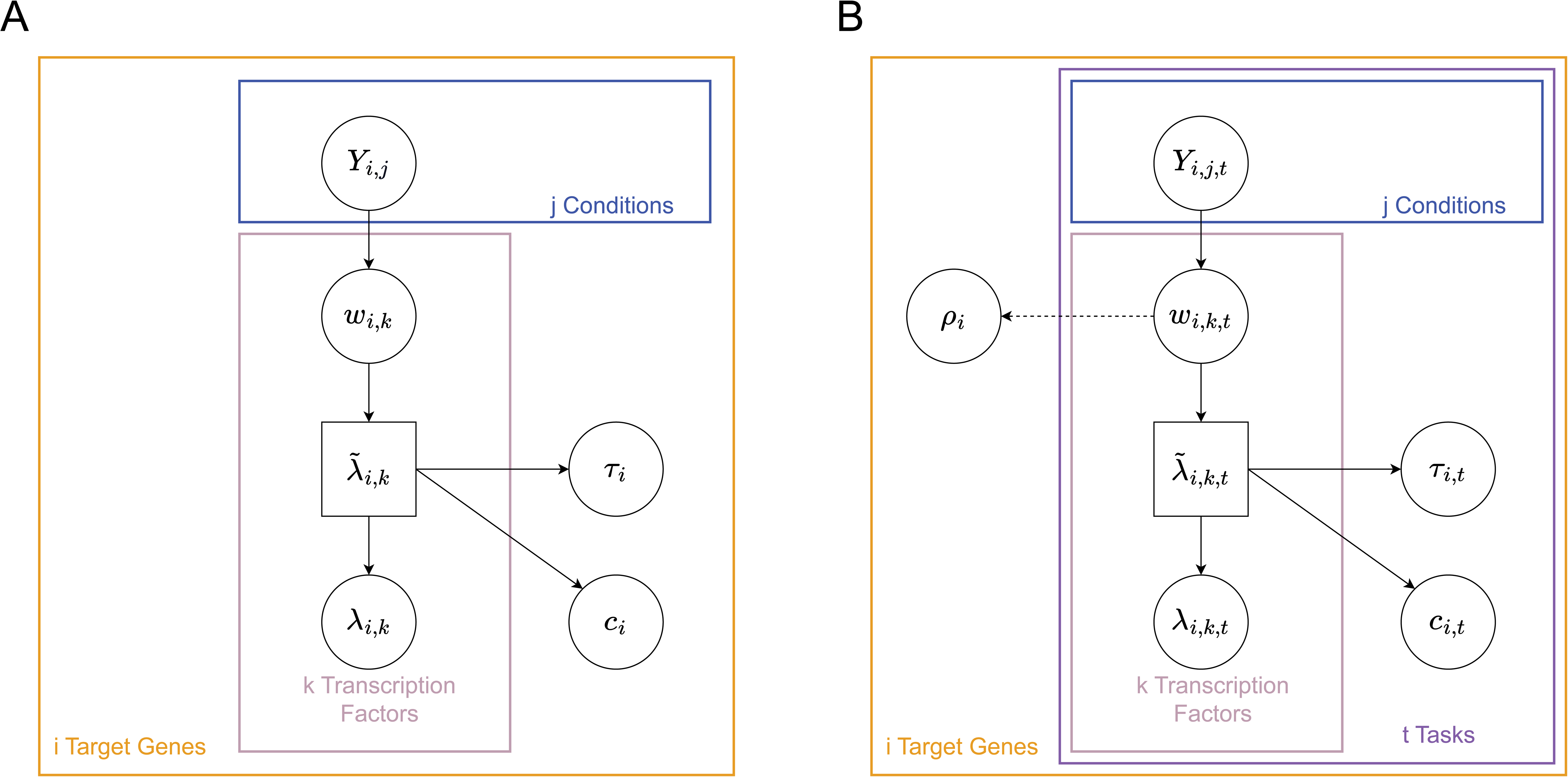
Graphical Bayesian Models Overview of variable dependence in GRN-BMuSeR in single task (A) and multitask (B) modes. Circular nodes correspond to sampled parameters and square nodes correspond to transformed parameters. Large colored boxes represent collections of variables that are sampled or transformed for each instance of the applicable subscript (*i* target genes in orange, *j* conditions in blue, *k* TFs in rose, *t* tasks in purple). Arrows represent dependence structure. In (B), the dashed arrow represents the addition of the learned correlation parameter which is only present in the GRN-BMuSeR Ext model. For detailed explanations of the variables in (A) see eqs. (2-6) and (B) see eqs. (8-12).

#### 0.6.6 Hyperparameters

The single-task and multitask models introduce three hyperparameters that must be chosen by the experimenter, slab degrees of freedom (*ν*), slab scale (*s*), and *P*_0_, with the extended model introducing an additional hyperparameter, *η*. As described in [20], *ν* and *s* correspond to a Student’s-t slab that regularizes the largest weights. As described in prior work, values for these two hyperparameters should be chosen such that the slab roughly corresponds to the size of the largest weights, but the exact value does not significantly change inference results as long as hyperparameters chosen do not produce a tight slab. [20]. For this reason, we fixed *ν* to 100 and *s* to 2, as this produced a large slab to avoid heavy influence on our results. *P*_0_ describes the predicted number of nonzero weights for each task and is also a component of the regularized horseshoe prior [20]. This value functions roughly as a budget for nonzero weights, and like the previous hyperparameter, inference is not sensitive to the exact value so long as the value is higher than the anticipated number of weights. For this work, we fixed *P*_0_ to 10, to encode prior information about the sparsity of gene-TF interactions [60]. *η* describes the shape of the *LKJ* prior placed on the correlation matrix in the extended model. We fix it to 1 for a uniform prior across possible correlation matrices to express our ignorance to the true correlation structure between the regulatory network underlying any given pair of orthologous genes. Since a correlation matrix is inferred for every gene, in theory this hyperparameter could be adjusted to account for prior knowledge in underlying network structure in a future work.

#### 0.6.7 Variable Selection and Interaction Weight Thresholding

The output of fitting the Bayesian model is a set of distributions of posterior weights *β_i,k_*. Since the horseshoe prior shrinks weights towards 0, but does not set the weights to 0, a variable selection procedure was required to select the weights that most likely correspond to true interactions. A “leave one out” (LOO) approach was used to capture weights describing both strong and weak but well-defined interactions. We first calculated the posterior mean of each weight distribution *β*^^^*_i,k_*. For the 10 highest values of *β*^^^*_i,k_*, we calculated a confidence score *c_i,k_*as previously described [61]. This confidence score measures the relative importance of a given transcription factor in explaining the resultant expression of a gene by taking the ratio of the performance of a full model versus a model from which that transcription factor has been removed. This ratio is proportional to the decrease in model performance caused by removing that TF from the model. We then tabulated the number of interaction edges remaining in the network that were greater than or equal to a score cutoff threshold in 0.01 increments. Orphan genes and TFs no longer connected at each threshold were removed. The elbow of the plot of sliding window score cutoffs vs the ratio of remaining edges to genes was taken as the variable selection threshold.

### 0.7 Biological analysis of GRN output

Resultant thresholded networks were analyzed in Cytoscape v3.10.4 using the “Analyze network” tool [62]. Power law plotting, fitting, and statistical analysis of the out-degree distributions was conducted using the poweRlaw package in R (v1.0.0 [63]; Figure S1). TF neighborhood analysis for each TF in the resultant GRNs was conducted by selecting the genes regulated by that TF together with genes regulated by the TF immediately upstream and the TF downstream of the TF of interest. The significance of enrichment in archaeal Clusters of Orthologous Genes (arCOG) [58] of gene sets within each TF neighborhood was calculated using a Fisher’s exact test. P-values of significance were adjusted for multiple hypothesis testing within each TF by false discovery rate using the Benjamini-Hochberg method. Global GRNs and subnetworks shown in the figures were visualized using Cytoscape.

#### 0.7.1 Implementation and Hosting

Our method is available as a python script and hosted on github at https://github.com/andrew-soborowski/GRN BMuSeR. Primary implementation of the model is done in Python while the Bayesian model is fit using a Hamiltonian Markov Chain Monte Carlo (HMCMC) approach in STAN, using the NUTs sampler [64]. As individual genes are inferred independently, parallelization is available for deployment on a high performance compute cluster. Sample scripts are provided for implementation using SLURM.

## Results

### 0.8 A Bayesian multitask model for joint inference of gene regulatory networks across species

In order to improve regulatory network inference by incorporating information across species, we developed a hierarchical Bayesian approach, GRN-BMuSeR (Gene Regulatory Networks from Bayesian MUlti-SpEcies Regression). This method is capable of sharing information between species while remaining robust when transferable information between species is unavailable. Our model takes in gene expression datasets for each species and outputs a rank-ordered list of confidence scores based on posterior strength of the inferred interaction for each species (Fig 2). Additionally, our model links information between each species on the basis of homologous genes and TFs between each species (See Methods).

**Fig 2.**
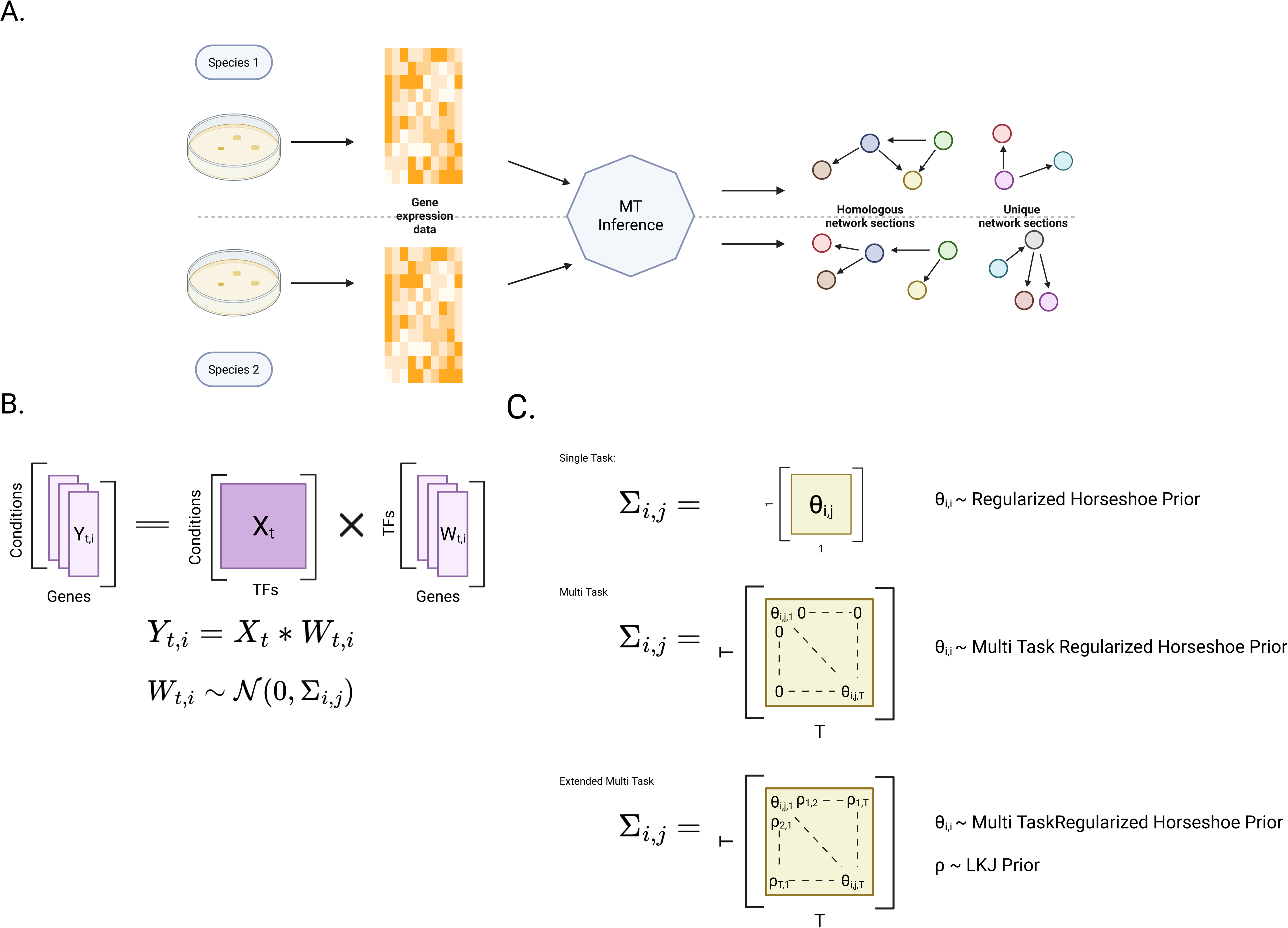
GRN-BMuSeR Overview (A) Overview of information flow for network inference. Expression data is collected from each species and processed, multitask inference is performed, and a network is generated for each species. Output networks will have homlogous sections for orthologous genes and TFs while allowing for unique sections of the network when orthology does not exist or function is not conserved. (B) Core regression modeling step. The expression of each gene in each condition is modeled as the weighted sum of each TF in each condition. A normal prior is placed on each weight term with mean 0 and variance/covariance Σ. (C) Definition of Σ depending on the type of inference performed. In single-task, Σ corresponds to a weight specific variance. In the multitask cases, Σ corresponds to a covariance matrix across orthologous TFs. In the extended case, task similarity is also modeled as covariance, whereas in the standard case all covariance terms are set to 0.

For each target gene-TF pair, we assumed that a direct regulatory link is the most likely feature to be conserved between species. Thus, we expected that our model would yield increased performance in species for which less data are available (referred to henceforth as “data poor species”) by transferring information on these inferred interactions between the species of interest. To map these links, our approach requires mapping node (genes and TFs) orthology. We created a one-to-one homolog mapping on the basis of reciprocal BLAST best hits (see Methods).

The core of our inference approach is implemented as a Bayesian hierarchical regression model, where we aim to predict resultant gene expression as a linear function of TF expression (Fig 2A). We learn weight terms, summarized as the mean of the posterior distribution over the weights, that link the expression of each TF to the expression of the target gene. We interpret a non-zero weight term as a prediction of regulatory interaction between a given TF and the target gene.

Since genes are typically regulated by a small number of TFs relative to the total number of available TFs [60, 65], we assume that our model should be sparse such that true weights of interactions between a given gene and most TFs would be zero. To account for this assumption of sparsity, we placed a regularized horseshoe prior on the variance of the weight terms [20]. This prior shrinks the variance of most of the weight terms to near zero, constraining their inferred value to in turn be near zero, implying no regulatory interaction between gene and TF. A small number of TFs with strong evidence escape this shrinkage and are only marginally reduced to improve stability in the sampling algorithm. To incorporate multitask information, we extended the regularized horseshoe prior by imposing dependence between the variances of the weights of homologous TFs (Fig 2B and C). Evidence of a regulatory interaction in one species that is conserved in the other species allows the variance term to escape shrinkage more easily in the second species. However, the inferred weight terms are conditionally independent such that some evidence of interaction in the second species is still required, albeit at a reduced level, for a non-zero weight term to be inferred.

Previous GRN inference algorithms have proposed using TF activity (TFA) as an alternative to TF expression as the predictive variable in regression [61, 66]. TFA is an abstraction that represents the ability of a TF to drive gene expression under a given condition. Incorporating TFA significantly increases performance as it can account for factors not included in raw expression data such as post-transcriptional regulation, post-translational modifications, and other molecular mechanisms that change TF binding affinity or behavior [61]. However, calculation of TFA values requires additional data beyond gene expression measurements, so its use is typically restricted to well-studied organisms such as *Saccharomyces cerevisiae* or *E. coli* [7, 67–69]. GRN-BMuSeR is capable of incorporating TFA when available to improve performance, but is also able to generate a network solely from TF expression data, as is common for applications involving understudied, data-poor organisms [4].

Additionally, we propose an extended model, GRN-BMuSeR Ext, in which we explicitly model correlation between the species. In this case, we impose a covariance matrix on the weight terms and jointly sample them. The variance terms are inferred as before through the regularized multitask horseshoe, but this time a flat *LKJ* prior is placed on the correlation at the gene level. This model allows for stronger information sharing between species under the assumption that relative effect sizes are similar across species.

### 0.9 Algorithm benchmarking

#### 0.9.1 GRN-BMuSeR Successfully Reconstructs Networks from Biological Data

To determine the performance of GRN-BMuSeR on experimental data, we benchmarked our approach against the Inferelator AMuSR (which we will refer to as simply Inferelator), an established multitask GRN inference algorithm [16]. The choice to benchmark against the Inferelator was guided by: (a) no equivalent algorithms exist to our knowledge; (b) the Inferelator is the closest match to our algorithm, although it is designed to incorporate multiple tasks from the same species in which every target gene and TF is represented in every task. Notably, for GRN-BMuSeR, this represents a special case, as the algorithm does not require the assumption that TFs and target genes are completely identical across tasks. We used transcriptomics datasets collected from two separate strains of the well-studied bacterium *Bacillus subtilis* exposed to a variety of genetic and environmental perturbations that were previously used to benchmark the Inferelator [16]. These datasets contain 429 and 269 transcriptome profiles for each strain, respectively. In addition, a high quality gold standard consisting of 3,040 experimentally validated interactions (153 TFs to 1,822 target genes) is available for this organism, making it a good choice to evaluate network edge recovery [16].

To evaluate the performance of GRN-BMuSeR for learning a GRN from these data, we inferred networks in three different ways. First, to confirm that GRN-BMuSeR is able to leverage cross-task information, we replicated a case study from the Inferelator’s publication that examines the interaction between target gene *ydfP* and TF *sigB* [16]. This interaction is well represented in task 2 but is difficult to infer in task 1 due to the different experimental conditions captured in each dataset [16]. For this case study, we used the entire gold standard in network generation to ensure that the maximum amount of information from the input datasets was incorporated. We generated networks using both the GRN-BMuSeR ST and GRN-BMuSeR models to observe the change in edge recovery given multitask information. In terms of variance explained (Fig 3A) and posterior weight (Fig 3B), the interaction between *ydfP* and *sigB* is well supported in task 2 for both resultant GRN-BMuSeR networks. Also as expected, recovery for the interaction in task 1 is poor in the single task network. Posterior weight was low enough that a value for variance explained is not calculated and defaults to zero. In the multitask network, the posterior weight of the interaction is recovered with the inclusion of the information in task 2, while variance explained remains low, but non-zero. These results demonstrate the ability of GRN-BMuSeR to incorporate multitask information successfully, as posterior weight is calculated using the evidence present in both tasks, while variance explained estimates the performance of interactions with high posterior weight uniquely in each task.

**Fig 3.**
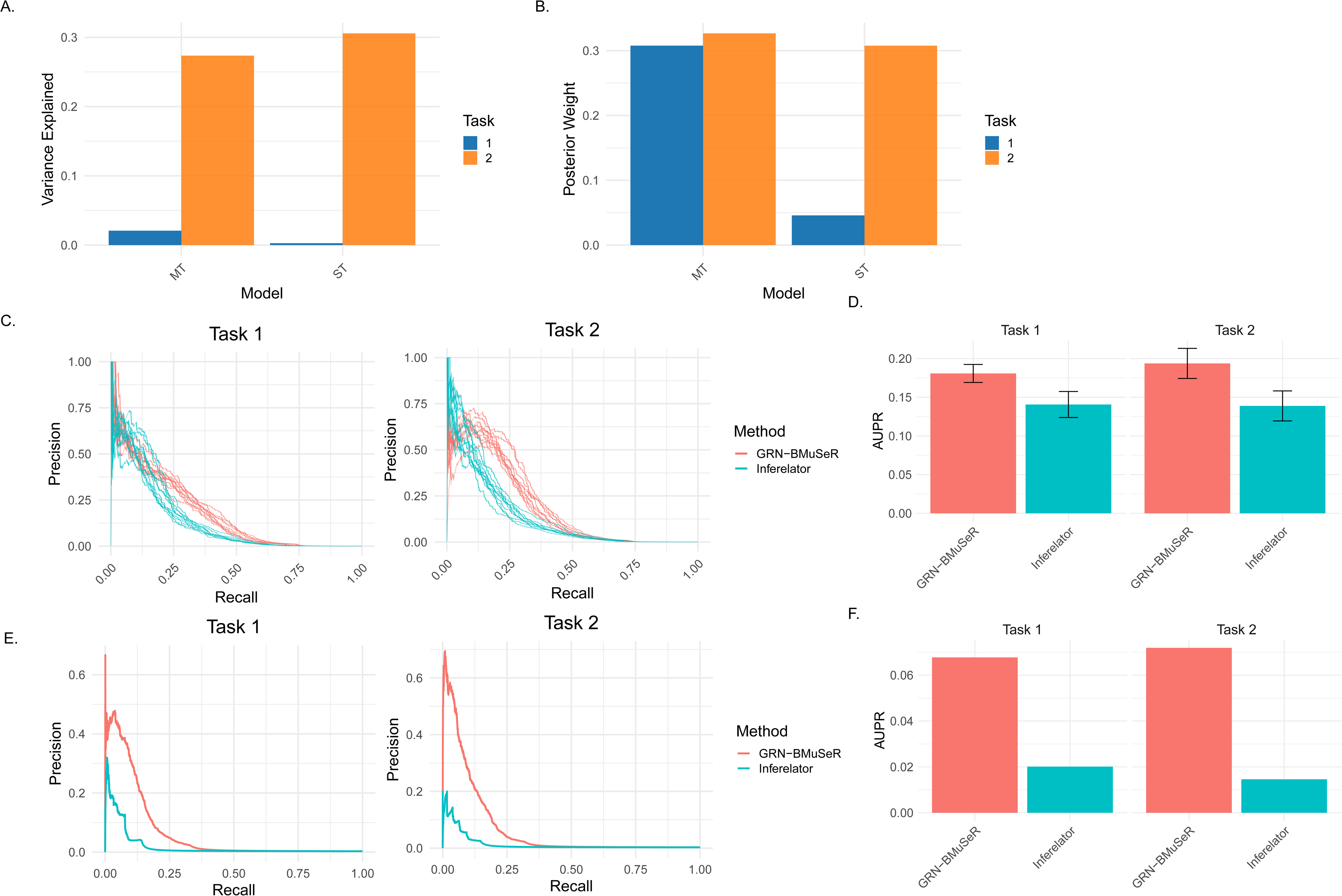
Benchmarking performance on *B. subtilis*. (A) Performance of GRN-BMuSeR in recovering a known interaction between *ydfP* and *sigB* using metrics of posterior weight or (B) variance explained. (C) Comparison of GRN-BMuSeR and Inferelator performance on the *B. subtilis* network using a prior. The prior was generated by sub-sampling 80% of the gold standard and the procedure was repeated 10 times. (D) Comparison of performance plotted in (C), expressed as AUPR. (E) As in (C) but comparing performance when no prior is available. As the gold standard was not subset, evaluation was performed once using the full gold standard. (F) As in (D) but comparing performance with no prior.

Second, we evaluated how GRN-BMuSeR performed in predicting all TF-gene interactions for *B. subtilis*. We inferred multitask networks using GRN-BMuSeR vs Inferelator using 80% of the gold standard interactions to build a TF activity matrix as the prior. The remaining 20% of interactions were held out randomly, repeating the inference procedure 10 times in order to average across the differential problem difficulty that arises from sampling. We evaluate performance using area under the precision-recall curve (AUPR) in order to measure how well the model recovers true positive interactions while avoiding false positives. We also expect that this metric will be useful in the context of an overwhelming number of true negatives in the network and an unknown number of false negatives in the gold standard, which are common in network inference [70]. We find that GRN-BMuSeR performs marginally better than Inferelator, though within the bounds of error (Fig 3C and D). Interestingly, the increased performance of GRN-BMuSeR appears to stem from a longer tail on the precision-recall curve which may suggest a trade-off between the recovery of less supported true-positive interactions at the cost of the inclusion of high confidence false-positives. This AUPR pattern may suggest that downstream examination of the resulting network would benefit from evaluating clusters of interactions around TFs, while placing less trust in any single predicted interaction. These results demonstrate the ability GRN-BMuSeR to perform in ideal conditions, where high quality orthogonal data are available to aid inference.

In the third and final benchmarking test, we compared our algorithm to Inferelator using data for which no prior is available, using raw TF expression as a predictor instead. Our approach outperforms the Inferelator in this case (Fig 3E and Fig 3F).

Taken together, these results suggest that our algorithm successfully generates network models from an established biological dataset, performing competitively with an existing method. Importantly, the improved performance on data lacking a TFA matrix prior suggests our approach is expected to perform well at network inference in understudied organisms for which TFA data are not available.

#### 0.9.2 GRN-BMuSeR remains robust to simulated species divergence

Although our method performs competitively to the Inferelator when inferring *B. subtilis* networks across strains, this dataset represents a special case for GRN-BMuSeR in which genes and TFs are identical between tasks. Therefore, we next evaluated the full capacity of our algorithm to infer GRNs in a multi-species setting in simulations where node identity and edge interactions progressively diverge between tasks (See Methods, Supplementary File 1). Briefly, we started by stimulating an underlying network structure for a single species, followed by a speciation event that separates the original into two descendant networks. In each descendant network (representing a species), we severed TF-target gene interactions (edges), removed target genes and TFs (nodes), introduced new nodes, and formed new edges. These steps were intended to simulate gene loss, gain, and overall network rewiring that naturally occurs in biological systems over evolutionary time. Each of these steps was done according to a metric we call “divergence”, which models the overall network differentiation in the system. When divergence=0, the underlying networks are identical. As divergence rises, the amount of gene turnover and network rewiring in each species increases proportionally. Next, given the underlying species networks, we generated mock experimental datasets through network perturbations. These datasets were then used as input for network inference, allowing for comparison against the known underlying network topology.

We used this approach to evaluate how our algorithm performed as tasks became increasingly dissimilar. To test this, we simulated 20 networks at each tested level of divergence, asking each of the 3 modes of GRN-BMuSeR to reconstruct them. We also reconstructed each network using linear regression as a baseline model (Fig 4A). Next, we compared each multitask model against the single-task GRN-BMuSeR ST model to evaluate performance differences specifically due to the incorporation of multitask information (Fig 4B). From these results, we observed an increase in performance using a multitask approach when the underlying network divergence was low. As task similarity decreased, the performance boost from multitask inference also decreased until an intermediate divergence level (~20). At divergence levels above 20 (very low network similarity across tasks), multitask inference became a performance liability (Fig 4). Additionally, we observed that GRN-BMuSeR Ext may perform slightly better than the base multitask model at low levels of divergence (Fig 4B). However, any performance gain was slight and not statistically significant given the small number of replicates sampled at divergence=0 (Welch’t T-Test, P-value = 0.39). Because of this marginal gain and the increased model complexity, we proceeded with GRN-BMuSeR as our primary model for subsequent analyses, though GRN-BMuSeR Ext may be useful on smaller subnetworks or other related problems. We concluded that performance improvements are directly proportional to the similarity between the underlying gene network structures of the species considered. Taken together, these results support our method’s ability to integrate multi-species information.

**Fig 4.**
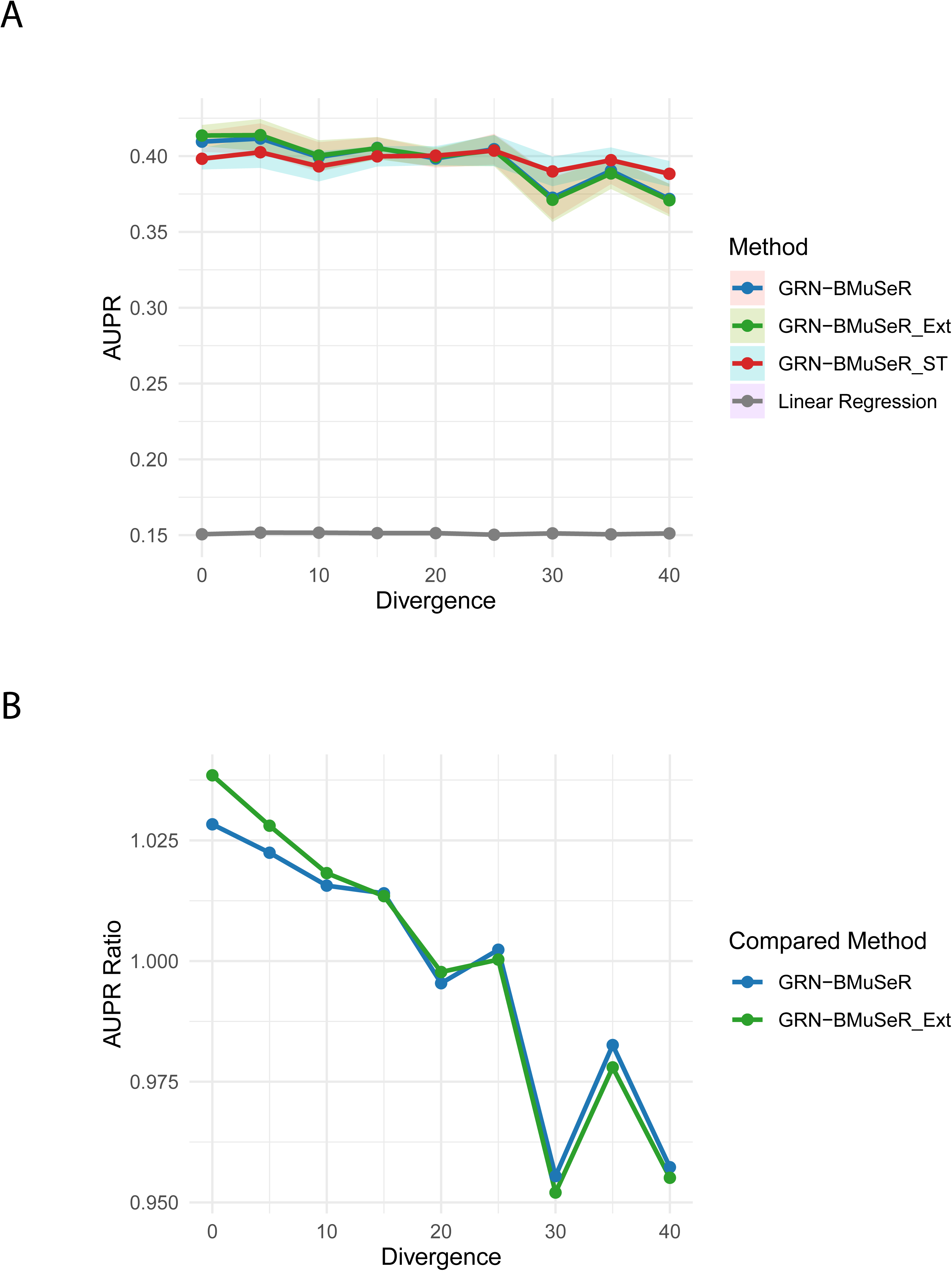
Benchmarking performance on Simulated Data. (A) Performance of GRN-BMuSeR as compared to its extended and single task versions on simulated data. Divergence represents the degree of separation between the underlying networks of the two tasks. AUPR is averaged across both tasks and the shaded regions represent standard error. Linear regression is included as a baseline level of performance. (B) Ratio in performance between the multitask versions of GRN-BMuSeR and the single task version. Each line represents a multitask approach divided by the single-task approach. Underlying data is the same as in (A).

In summary, taking benchmarking results with experimental and simulated data together, we demonstrate that GRN-BMuSeR performs comparably with established approaches for inferring a GRN on a biological dataset from a well-studied bacterial species while remaining robust to differences in underlying network structure.

### 0.10 GRN-BMuSeR jointly infers GRNs across species of understudied hypersaline adapted archaea

#### 0.10.1 Generation of a transcriptomics compendium for *Hfx. volcanii*

While a large expression dataset is already publicly available for *Hbt. salinarum* [40], transcriptomics experiments for *Hfx. volcanii* are limited and fragmented, motivating the compilation of a unified dataset to enable joint network inference and broad community use. Using RNA-seq, we generated 149 new transcriptome profiles across five different media conditions and 34 knockout strains (Fig 5A, Supplementary Tables 1 and 2). We combined this with previously published, publicly available extant data from the research community and our lab for a total of 282 transcriptome samples representing 40 strains grown under 13 different environmental conditions, a dataset size on par with those used historically for the first GRN inference models for individual species of archaea [4, 10]. According to correlation analysis, samples were reproducible within experiments between replicates of the *Hfx. volcanii* dataset (Fig 5A). Principal component analysis demonstrated that 12 PCs were required to explain 80% of the variance, suggesting high dataset complexity. At the same time, variance was saturated within the dataset (96% variance explained by 55 cumulative PCs), implying that the dataset captured a sufficient diversity of transcriptome activity. To determine whether this variance was attributable to biological effects, we conducted variance partitioning analysis using a linear mixed model as described in [57] (See Methods). Growth medium treatment (“condition” variable) explains a median of 30% of the variation broadly distributed across genes (Fig 5C). In contrast, lab of origin explains a significantly lower proportion of expression variance (Paired T-test, mean difference 0.054, *p <* 2.2 × 10*^−^*^16^) that disproportionately affects a small number of genes (i.e. the “lab” variable explains zero variance for ~ 500 genes). Thus, although there is some nesting of variance of media condition within the “lab of origin” variable, a higher proportion of variance is explained by growth medium than lab. This result suggests that important biological variation has been captured because a large proportion of the transcriptome program is known to be affected by variation in nutritional regimes in these species [71, 72]. A median of 28% is explained by unknown factors (residual fraction). Together, these analyses suggest that the *Hfx. volcanii* expression compendium captures substantial meaningful biological transcriptome variation. This compendium substantially increases the number of the transcriptome datasets available for *Hfx. volcanii*.

**Fig 5.**
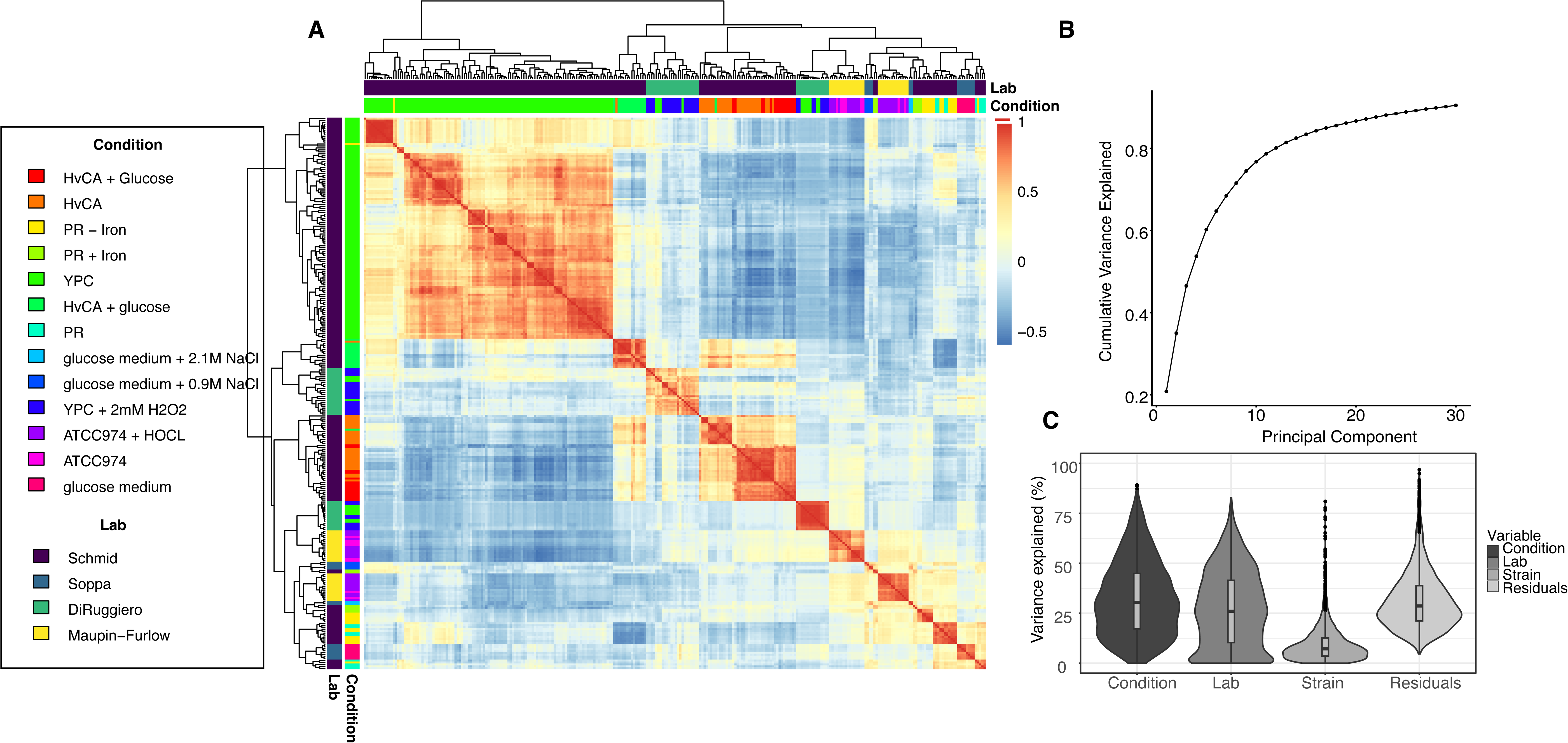
A large expression compendium for *Hfx.* volcanii captures transcriptome variation. (A) All-by-all correlation heatmap comparing transcriptome profiles. Each square of the heatmap represents one transcriptome sample. Equivalent samples are shown along the diagonal and the color scale (see legend at right) for the off-diagonal represents the Spearman correlation coefficient. Color bars above and left of the heatmap represent condition and lab according to the legend at left. Dendrogram represents hierarchical clustering across samples (see Methods). (B) Cumulative variance explained (y-axis, given as a fraction from 0 to 1) for each additional eigenvector (x-axis) of the principal component analysis (PCA) of the data. The first 30 components are shown for clarity. (C) Violin plot showing the distributions of percent variance explained (y-axis) by each treatment variable (x-axis). Residual variance (far right) represents unexplained variation not captured by other declared variables.

#### 0.10.2 Joint inference and variable selection thresholding yields high-confidence global GRNs for two species of haloarchaea

We applied this dataset and that from *Hbt. salinarum* [40] to jointly learn GRNs de novo across *Hbt. salinarum* and *Hfx. volcanii* using GRN-BMuSeR. Given the limitation that gold-standard regulatory interactions are scarce relative to those in well-studied organisms, we used the priorless approach as described in the *B. subtilis* benchmarking case above (Fig 3E and F, see also Methods). The generated raw networks comprised scored predictions, or weights, for every possible interaction between the input TFs and target genes (for TF definition, see Methods). Variable selection was then conducted post-hoc to maximize high-confidence, biologically meaningful predictions and ensure network sparsity (Methods, Figure 6A and B). Briefly, we generated confidence scores for each interaction based on posterior weight means (“leave one out”, or LOO, approach, see Methods). Subsequently, species-specific network thresholds were calculated from the elbow of a plot of LOO confidence scores vs the ratio of total interactions to the remaining incoming interactions per gene. This threshold was chosen to balance strong explanatory power (interactions with a large confidence score) with network sparsity (progressive elimination of interactions). Applying the threshold resulted in a score cutoff of 0.09 in *Hbt. salinarum* (Fig 6A) and 0.19 in *Hfx. volcanii* (Fig 6B).

**Fig 6.**
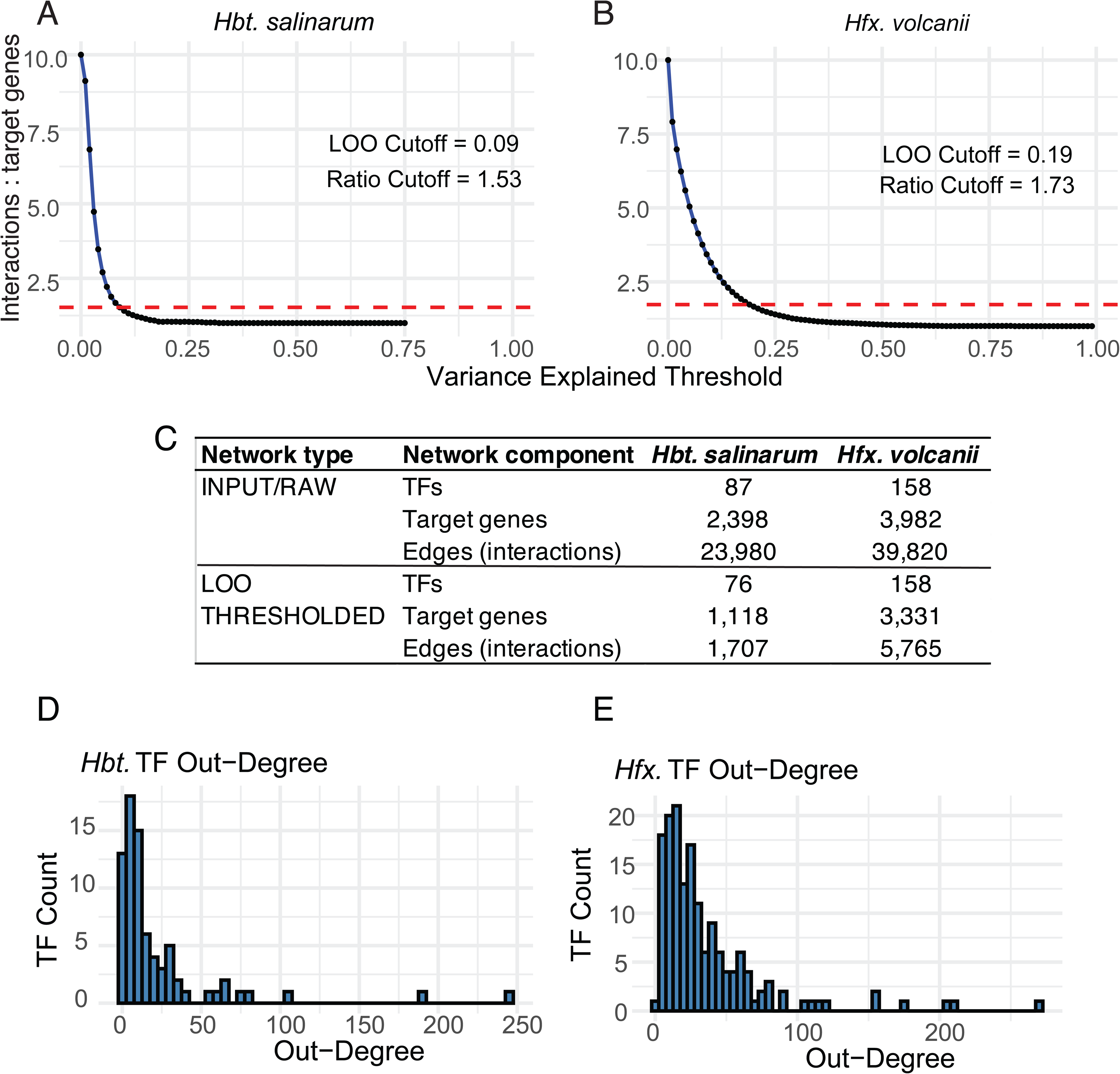
Network variable selection and resultant global network features. (A) For *Hbt. salinarum*, a sliding window of the variable selection leave one out (LOO) threshold score threshold (x-axis) is plotted against the ratio of total network interaction edges to the total remaining target genes (y-axis). The dotted red line indicates the elbow of the plot and therefore the chosen score threshold. LOO score threshold and y-axis ratio cutoffs are written in the plot insets. (B) As in (A) but for *Hfx. volcanii*. (C) Table of network edge and node enumeration. Raw network output is given above, LOO thresholded network below. Numbers are also described in the text. (D) Distribution of out-degrees for target genes in *Hbt. salinarum*. (E) Distribution of out-degrees for TFs in *Hfx. volcanii*.

We analyzed the global features of the resultant thresholded networks to determine their capacity for providing biological insights. The *Hbt. salinarum* network contains 1,707 interactions between 1,158 targets and 76 TFs out of 2,647 total protein-coding genes encoded in the genome (Supp Tables 3 and 4, Fig 6C). The *Hfx. volcanii* network was slightly larger given the greater number of protein-coding genes encoded in its genome: 5,765 interactions across 3,331 targets and 158 TFs from the initial input of 3,982 targets and 158 TFs (Fig 6C). The network density, a proxy for sparsity, for both species was low (0.003 for *Hbt. salinarum* and 0.001 for *Hfx. volcanii*). The low average number of node neighbors also suggested sparsity (2.98 and 3.41 for *Hbt* and *Hfx*, respectively). The connectivity of the network nodes for each species, or degree distribution, was heavily skewed with a long right-handed tail, suggesting the presence of network communication “hubs” (Fig 6D and E, Fig S1B and D, [73]). For example, the average out-degree for TFs in the *Hbt. salinarum* was 25.5 (9 global TFs with out-degree *>* 50; 60% *<* 30; Fig 6D). For *Hfx. volcanii*, the average out-degree across all TFs is 36.47, (33 global TFs with out-degree *>* 50, 79% TFs *<* 30; Fig 6E). The high out-degree TFs are predicted to be network hubs, which can be interpreted as putative global regulators, whereas the majority of TFs are predicted to function as specific regulators. In-degree distributions were also right-skewed in both species; however, the inference procedure constrains the in-degree so this was a less informative metric here (Fig S1A and C). The exponent of the power law fit to the log-log tail of each out-degree distribution (gamma) was *>* 1, suggesting that each network was approximately scale-free (Fig S1D, [74], methods). Taken together, these global network metrics are well aligned with the small-world hub-and-spoke structure and sparsity criteria expected for biological networks [73–75]. This result suggests that the inferred networks may enable useful biological predictions of gene regulation.

#### 0.10.3 GRN-BMuSeR inferred networks recapitulate known functions and predict expanded functions for well studied TFs

To explore the biological predictions, we sought to demonstrate that the inferred GRNs can recover previously known information about well studied TFs. We examined the functions of genes in the neighborhood of each TF, which included target genes both one and two degrees removed from each TF, inclusive of incoming and outgoing edges (Fig 7A, Fig 8A and B). This approach enabled us to circumvent the caveat that GRN inference procedures do not distinguish direct from indirect interactions [9]. For gene sets within each TF neighborhood, we calculated the significance of functional enrichment in archaeal Clusters of Orthologous Genes (arCOG) categories (Fig 7B and C, [58]). In *Hbt. salinarum*, 33 of 76 TFs (43%) were predicted to regulate gene neighborhoods with arCOG annotation enrichments (Fig 7B, Supplementary Table S4). These arCOG functions fell into 11 categories, six of which had metabolic functions.

**Fig 7.**
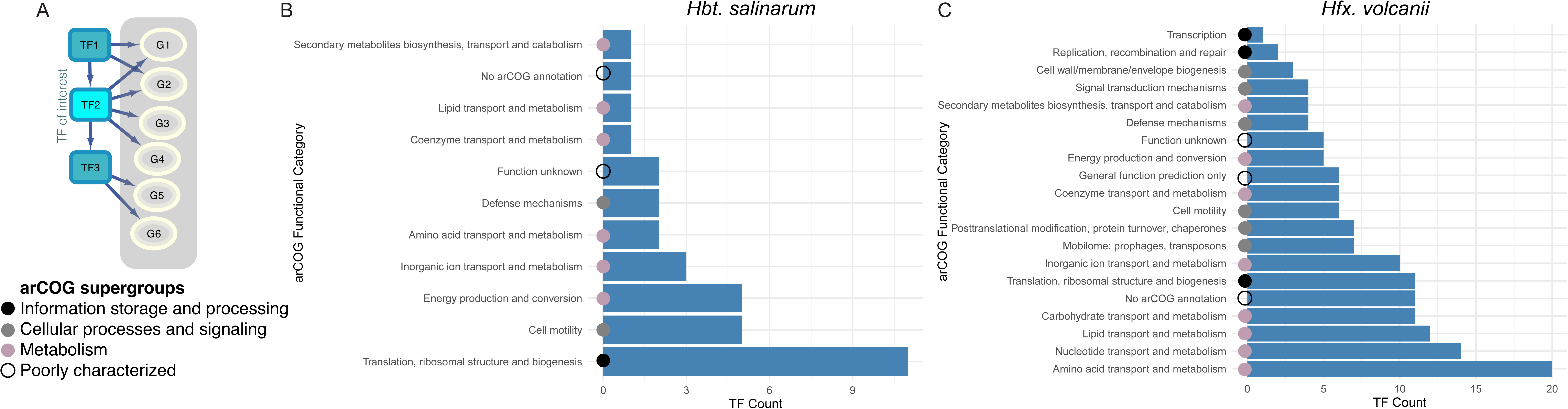
Global functional predictions for TF neighborhoods. (A) Diagram illustrating a TF neighborhood. The TF of interest (TF2) is highlighted in cyan, and its neighborhood includes upstream TFs (represented by “TF1” in the teal box), downstream TFs (“TF3”) and target genes of all three TFs (grey circles G1-G6). (B) Bar plots of significantly enriched functions in arCOG categories for *Hbt. salinarum*. X-axis represents the number of TFs whose neighborhoods are enriched for the functional categories given on the Y-axis. Colored dots at the base of each bar shows the membership of the arCOG category in each supergroup according to the color legend provided in the lower left corner of the figure. (C) Neighborhood gene enrichments for modeled TFs in *Hfx. volcanii*. Enrichment scores were calculated based on the functions of genes in each TF neighborhood. Enrichment analysis included all modeled TFs in each species. Enrichments were calculated using a Fischer’s exact test. P-values of significance were adjusted for multiple hypothesis testing by false discovery rate (FDR) (see Methods and Table S4).

**Fig 8.**
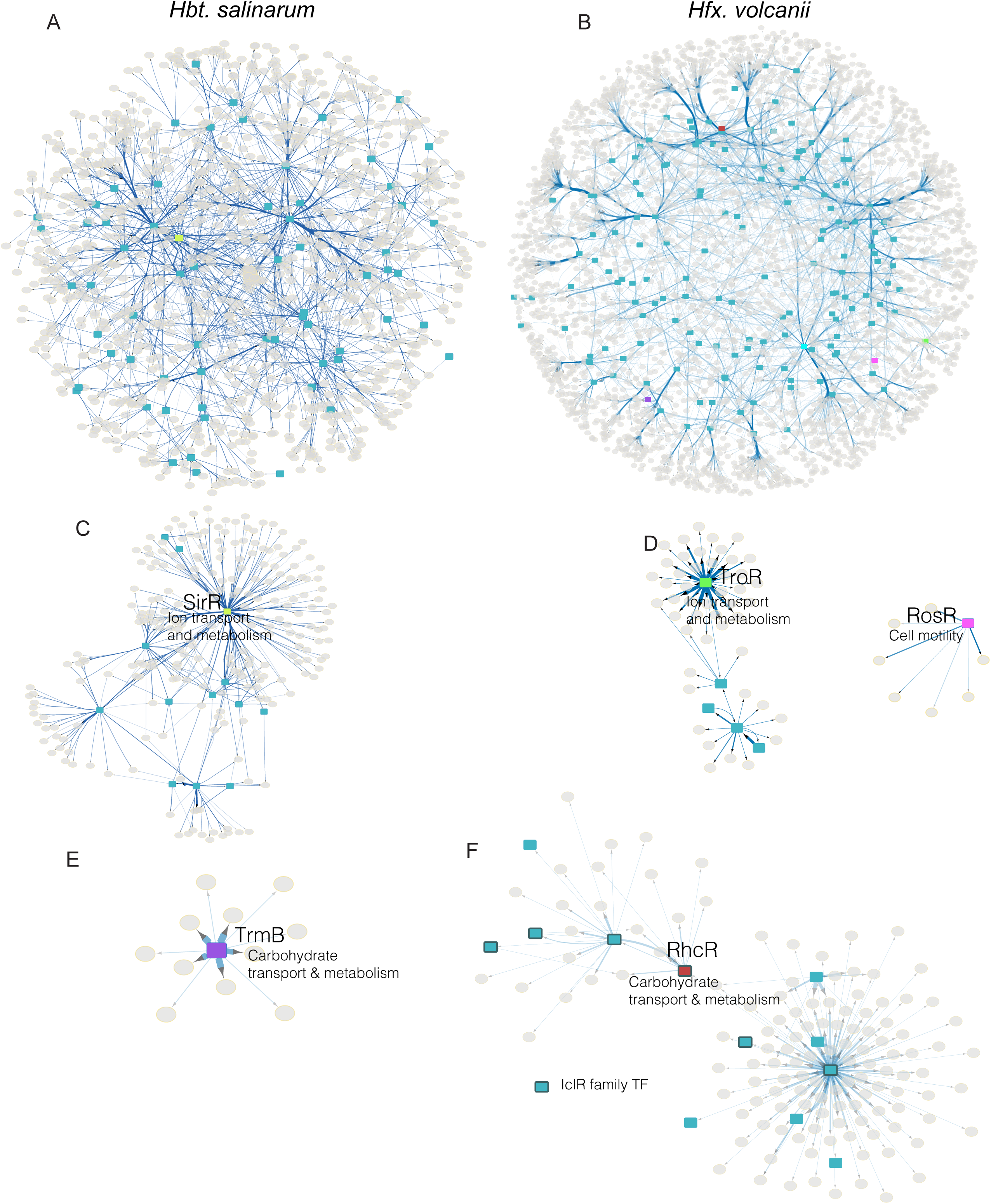
Inferred GRNs recapitulate known TF functions and generate hypotheses for novel TF functions. (A) Cytoscape global network visualization for the confidence thresholded GRN inference output in *Hbt. salinarum*. TFs are represented by blue rectangles and target genes by grey circles. TFs with other colors show the highlighted TFs in context, and correspond to the subnetworks below. Regulatory edges are shown as blue arrows. Edge thickness corresponds to confidence score strength and transparency corresponds to posterior mean (higher saturation represents higher, more confident weight). Orphan interactions not connected to the larger network are not shown for simplicity, but these are accessible in the Supplementary Table S5 containing all interactions, confidence scores, and posterior means for exploration in Cytoscape. Edges outgoing from TFs are bundled for clustered genes. (B) Cytoscape visualization for the *Hfx. volcanii* network. (C) SirR function in metal homeostasis regulation was recapitulated by the GRN inference predictions. Subnetwork visualization is shown for *Hbt. salinarum*. Chartreuse colored TF node highlights SirR in the corresponding node color in the global network in (A). (D) Subnetwork visualizations for *Hfx. volcanii* including TroR (bright green), RosR (pink). (E) Subnetwork visualization for *Hfx. volcanii* TrmB (purple). (F) Subnetwork visualization for RhcR (red). Across all panels, network layout was conducted using the edge weighted spring embedded method.

The majority of TF neighborhoods were predicted to function in translation (11 of 32 total enriched neighborhoods, Fig 7B). In contrast, the *Hfx. volcanii* GRN had a larger fraction of TF neighborhoods with functional enrichments in 20 arCOG categories (106 of 158 TFs, 67%, Fig 7C). Only four of these *Hfx. volcanii* TFs had poorly characterized or unknown functions (4.6%). As in *Hbt. salinarum*, most *Hfx. volcanii* neighborhoods were predicted to function in metabolism (8 of 20 category enrichments); however, a larger fraction of TFs in *Hfx. volcanii* were predicted to be involved in cellular processes and signaling than in *Hbt. salinarum*. More TFs in *Hfx. volcanii* were predicted to regulate genes of unknown function (16%) (Fig 7C). Taken together, these GRN-wide functional enrichments suggest that the model output can be used for hypothesis generation.

To further investigate these general functional insights, we asked whether the inferred GRNs were able to qualitatively recapitulate previously published biological functions for TFs. We compared TF neighborhood arCOG predictions for each TF to experimentally characterized TF functions reported in the scientific literature. For *Hfx. volcanii*, 22 TFs used for inference have experimentally characterized and published functions. The function of 18 of these TFs were predicted to regulate a neighborhood with an arCOG enriched function (Table S4). Of these, 11 TF functions were correctly predicted, *i.e.* these TFs were predicted to regulate a gene neighborhood with significant enrichment in at least one arCOG functional category (out of 26 total arCOG categories, [58]) that corresponded with the published TF function (Table 1, Fig 8B,D-F, Supplementary Figure S1E). For *Hbt. salinarum*, 13 of 22 known TFs were predicted to regulate neighborhoods with enriched functions (Table S4). Six of these TF functions were correctly predicted by GRN-BMuSeR (Table 2, Fig 8A and C, Supplementary Figure S1F). Strikingly, predictions for homologous TFs recapitulated known TF functions across species. For example, the metal-dependent DtxR protein family TF SirR in *Hbt. salinarum* is known to repress metal uptake under iron and manganese replete conditions [34, 76]. Under similar growth conditions, the DtxR family TF TroR in *Hfx. volcanii* represses iron uptake [44]. For these homologous TFs, the network inference procedure correctly predicted a gene neighborhood significantly enriched in the arCOG category “Inorganic ion transport and metabolism” (Fig 8C and D, Table S4). SirR is predicted to control 243 genes, including 11 other TFs (Fig 8C). TroR is predicted to control 63 target genes, including four other TFs (Fig 8D).

**Table 1.** Experimentally characterized, correctly predicted TF neighborhood arCOG enrichments in *Hfx*.

| NCBI gene ID | HVO ID | Common name | arCOG Functional Category | adjusted p-value | Citation |
| --- | --- | --- | --- | --- | --- |
| HVO_RS07290 | HVO_0538 | Idr | Inorganic ion transport and metabolism | 1.66E-08 | [44] |
| HVO_RS07885 | HVO_0662 | ThiR | Coenzyme transport and metabolism | 1.40E-03 | [82] |
| HVO_RS08205 | HVO_0730 | RosR | Cell motility | 3.61E-02 | [45] |
| HVO_RS08835 | HVO_0863 | TroR | Inorganic ion transport and metabolism | 6.26E-09 | [44] |
| HVO_RS15255 | HVO_2194 | Sir2 | Translation, ribosomal structure and biogenesis | 4.19E-04 | [83] |
| HVO_RS16130 | HVO_2374 | PhoU | Lipid transport and metabolism | 1.26E-04 | [83] |
| HVO_RS17680 | HVO_2688 | TrmB | Carbohydrate transport and metabolism | 4.87E-03 | [42] |
| HVO_RS19070 | HVO_2970 | OxsR | Coenzyme transport and metabolism | 1.06E-03 | [49] |
| HVO_RS00550 | HVO_B0114 | RhcR | Carbohydrate transport and metabolism | 0.0132 | [78] |

**Table 2.**
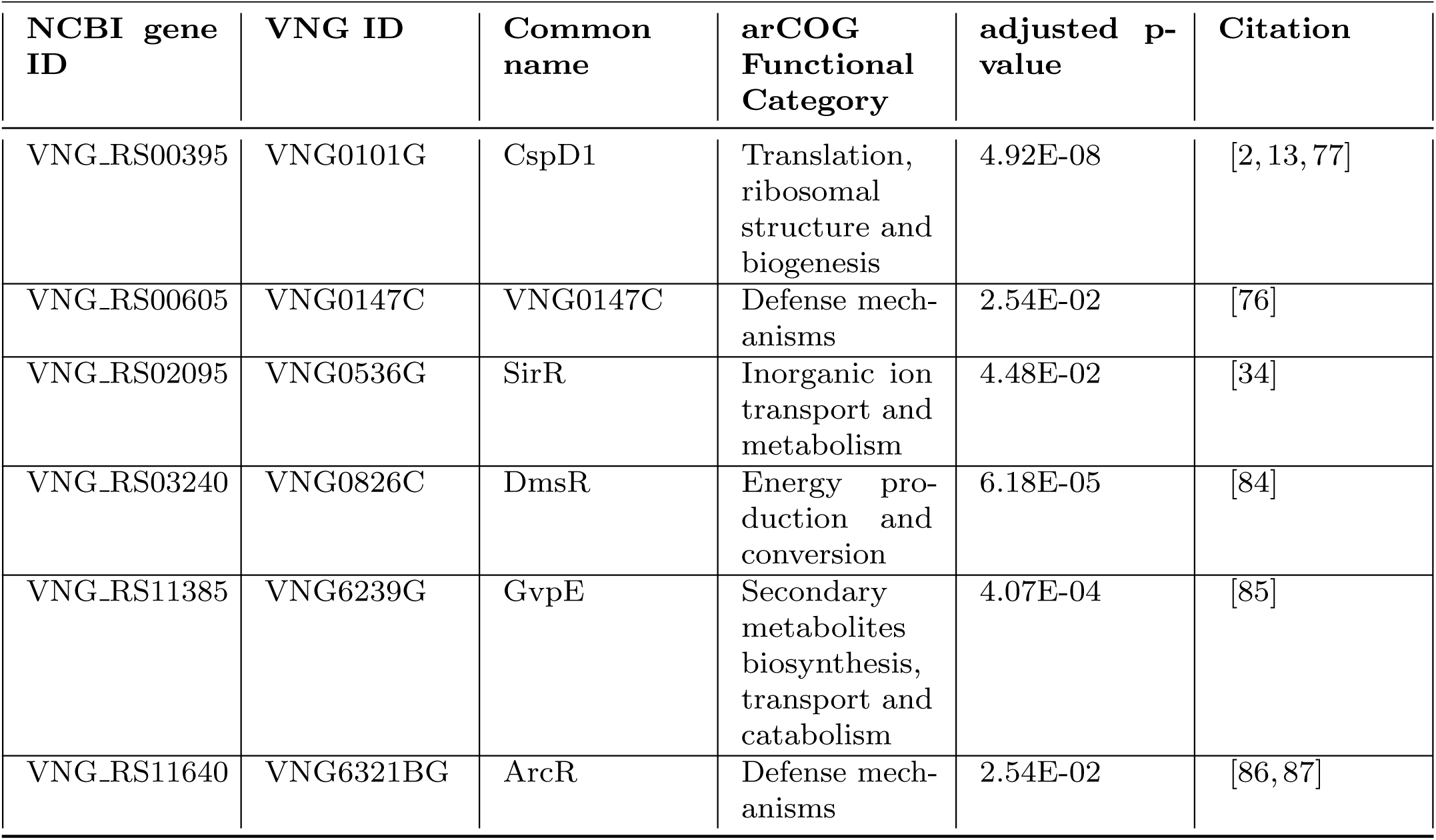
Experimentally characterized, correctly predicted TF neighborhood arCOG enrichments in *Hbt*.

| NCBI gene ID | VNG ID | Common name | arCOG Functional Category | adjusted p-value | Citation |
| --- | --- | --- | --- | --- | --- |
| VNG_RS00395 | VNG0101G | CspD1 | Translation, ribosomal structure and biogenesis | 4.92E-08 | [2, 13, 77] |
| VNG_RS00605 | VNG0147C | VNG0147C | Defense mechanisms | 2.54E-02 | [76] |
| VNG_RS02095 | VNG0536G | SirR | Inorganic ion transport and metabolism | 4.48E-02 | [34] |
| VNG_RS03240 | VNG0826C | DmsR | Energy production and conversion | 6.18E-05 | [84] |
| VNG_RS11385 | VNG6239G | GvpE | Secondary metabolites biosynthesis, transport and catabolism | 4.07E-04 | [85] |
| VNG_RS11640 | VNG6321BG | ArcR | Defense mechanisms | 2.54E-02 | [86, 87] |

Species-specific TF information was also recovered by GRN-BMuSeR. For example, RosR is a highly conserved archaeal-specific TF that has been demonstrated to be required for activation of genes that function in cell motility in *Hfx. volcanii* [45]. RosR function in *Hbt. salinarum* has been extensively rewired [45] to regulate oxidative stress response [2, 13, 77]. The GRN predicts a small regulon with a sole functional enrichment in cell motility genes in *Hfx. volcanii* (*p <* 0.04, Table 1, Fig 8D, right), recapitulating these previous findings. Although RosR in *Hbt. salinarum* did not have a functional enrichment (Table S4), known direct targets (e.g. heat shock genes such as *hsp20*) were recapitulated in this species (Table S5, VNG6201G, operon VNG1800H / 1801G). Given that most arCOG category enrichments were functions associated with metabolism (Fig 7), we visualized subnetwork diagrams for TF neighborhoods with these predicted functions. For example, in *Hfx. volcanii*, TFs TrmB and RhcR have been demonstrated to regulate the central carbon metabolic pathways gluconeogenesis and rhamnose metabolism, respectively [42, 78]. The neighborhood of each TF was significantly enriched in the functional class “carbohydrate transport and metabolism”, demonstrating that GRN-BMuSeR accurately predicts the known function of these TFs (Fig 8E and F, respectively). Taken together, these results demonstrate that our approach recovers known functional categories of target genes for specific TFs, suggesting that the GRN predictions represent biological insights.

#### 0.10.4 TF neighborhood enrichments generate novel predictions for understudied TFs

Given that the inference procedure was able to recapitulate known TF functions, we next sought to generate testable hypotheses about the functions of poorly characterized TFs in halophilic archaea. We applied the same TF neighborhood enrichment analysis to all transcription factors in LOO thresholded inferred networks for both species (Fig 8

A and B, Supplementary Tables 3 and 4). We first generated hypotheses starting with a functional category of interest to search for potentially implicated TFs. For example, the regulation of carbohydrate metabolism and energy conversion is of intense interest to the field because of the suitability of haloarchaeal enzymes and metabolic pathways to bioengineering applications [12]. TF neighborhood enrichments in the “carbohydrate transport and metabolism” category included one TF in *Hbt. salinarum* (VNG1903C, previously uncharacterized) and 15 for *Hfx. volcanii*, 13 of which remain unstudied (Table S4). Although VNG1903C is annotated in NCBI as a TrmB family TF, it bears weak sequence identity (29%) to the N-terminal DNA-binding domains of characterized TrmB metabolic regulators [42, 79]. No homology was detected in the C-terminal carbohydrate-binding domain or by structural alignment of predicted protein structures. This hypothesis demonstrates that GRN predictions are an essential part of TF functional discovery. For the “energy production and conversion” category, six TF neighborhoods were enriched in *Hbt.* (4 novel) and seven in *Hfx.* (all novel). To gain further insight into the predicted function of these TFs via “guilt by association”, we examined other TFs and genes in the neighborhoods of these novel TFs. For example, the rhamnose catabolism regulator RhcR shares a TF neighborhood with four other IclR family TFs, including the known xylose catabolic regulator XacR [80] (Fig 8F).

Across bacteria and archaea, IclR TFs have been implicated in nutritional regulation [78, 80, 81], so the three unknown IclR TFs in this neighborhood are strong candidates for metabolic regulators (HVO B0119, HVO A0527, HVO 2110, Table S3). Taken together, these predictions demonstrate the ability of our approach to generate hypotheses for future work on understudied TFs, either through a hypothesis about a functional category of interest or through a hypothesis about a specific TF for the functional categories and genes that it may regulate.

Taken together, these predictions demonstrate the ability of our approach to generate hypotheses for future investigation of understudied TFs, either by identifying TFs associated with a functional category of interest or by predicting the functional categories and genes regulated by a specific TF.

## Discussion

Here we present GRN-BMuSeR, a novel algorithm capable of inferring gene regulatory networks jointly between closely related species. Our approach leverages Bayesian linear regression in a hierarchical setting to provide a highly interpretable model in which TFs and target genes are directly linked via learned interaction weights (Fig 1 and 2). This model provides an advantage relative to previous multitask inference approaches in that interactions between both homologous and non-homologous genes are learned [16, 18]. This algorithm performs comparably to existing multitask procedures when TF activity and orthogonal data are used as priors, but demonstrates better performance when inferring based on TF expression data alone (Fig 3, [16]). Through simulations, we demonstrate the usefulness of the model for inference of closely related networks with non-identical nodes (Fig 4). We computationally define network divergence, a parameter intended to address a long-standing question of how GRN structures change as they evolve. Although previous algorithms have explicitly encoded phylogenetic information to inform GRN inference [88], our divergence parameter is agnostic to phylogeny. We demonstrate a fundamental trade-off between network structural relatedness and algorithm performance (Fig 4). The organisms included here in multitask inference, *Hfx. volcanii* and *Hbt. salinarum*, are related at the Order level, suggesting that multi-species inference is still effective at relatively large phylogenetic distances (~600 Mya [22, 23]). However, in cases with extreme network divergence, the single-task GRN-BMuSer ST version we present here can be used for single species inference for improved runtime and decreased model complexity (Fig 3). An important future research question will be to determine how biological network structural divergence scales with phylogenetic relationships across the tree of life.

Importantly, we show that jointly inferred GRNs for these hypersaline-adapted, extremely stress resistant archaea recapitulate known biology across both organisms (Fig 8, [34, 44, 78]). Model analysis provides experimentally testable hypotheses regarding novel TFs (Fig 8, Fig S1E and F). GRN-BMuSeR provides these predictions in the absence of a TF gold standard dataset by implementing a heuristic that balances the average number of incoming interactions per target gene with a sliding threshold on interaction explanatory power (Fig 6). Although previous iterations of Inferelator also conducted inference without a gold standard, the current model allows for joint multi-species inference [4].

In the long term, a fruitful research avenue would be to develop TFA priors for understudied species from orthogonal experimental datasets such as TF-DNA interactions (ChIP-seq) across all encoded TFs. Indeed, we observed substantially improved performance with benchmarking on *B. subtilis* data upon inclusion of TFA priors from a gold standard dataset (Fig 3) [16]. TFA is an abstraction of how and when a TF exerts direct regulatory influence on its target genes [7]. TFA inclusion therefore improves inference relative to raw expression data because it captures situations in which, for example, TF-DNA binding activity is regulated at the post-translational level. The current model and previous GRNs for archaea are similarly limited in this case: they detect TF-gene interactions for the ~ 50% of TF encoded genes that are regulated at the level of gene expression [4, 9]. Furthermore, while we were able to glean important biological insights from GRN BMuSeR output using arCOG functional enrichments of gene neighborhoods ([58], Fig 8, Table S3 and S4), continual improvements of genome annotations will enhance future understanding of TF functions, the target genes they regulate, and TFA priors. For example, structural predictions and homology searches using tools such as AlphaFold and FoldSeek can enable detection of homology in cases of low sequence-based homology [89–91]. Recent comprehensive literature syntheses and large-scale proteomics have improved genome annotations and related databases (e.g. NCBI, arCOG) for hypersaline adapted archaea in particular [92, 93].

After considering this full scope of the GRN model we created, we emphasize the biological utility of GRN BMuSeR. We present testable hypotheses for rapid experimental characterization of GRNs in multiple understudied but biotechnologically important organisms simultaneously [94]. It is important to note that there is no theoretical limit to how many species can be simultaneously jointly inferred using this method, provided their underlying networks are within the divergence bounds where multitask GRN-BMuSeR outperforms the single-task version (GRN-BMuSeR ST, Fig4). The two organisms jointly inferred here were related at the level of phylogenetic Order, suggesting that future work with GRN-BMuSeR could enable simultaneous inference of directed gene interaction networks for entire groups of understudied organisms. With the GRN-BMuSeR framework, we have laid important computational groundwork for future analyses, which includes (a) systematic assessment of uncertainty in GRN predictions, and (b) further extension of the additive linear model to incorporate more complex multiplicative effects such as combinatorial TF control of gene expression, among other myriad extensions. Taken together, we conclude that our model presents a biologically informative computational framework for joint multi-species GRN inference across genera.

**Fig 9. S1 Figure. Additional information about GRN inference output network analysis.**

(A) In-degree histogram for target genes in *Hbt. salinarum*. Associated statistics are given next to the figure. (B) Power law fit (red line) to the tail of the log-log TF out-degree distribution for *Hbt. salinarum*. Each dot represents a TF. (C) In-degree histogram for target genes in *Hfx. volcanii*. (D) Power law fit to the tail of the log-log TF out-degree distribution for *Hfx. volcanii*.

## Supporting information

**S1 Table.**

**Strain list:** Table of strains and RNA-seq experiments utilized in *Hfx. volcanii* for both data generated for this work and pre-existing data sources.

**S2 Table.**

**Normalized RNA-seq data:** Table of RNA-seq data from *Hfx. volcanii* normalized across strains.

**S3 Table.**

**TF annotations:** Annotations for all TFs used for input into the inference procedure

**S4 Table.**

**TF target gene neighborhood arCOG functional enrichments:** Table of enriched pathways among TF neighborhoods in *Hbt. salinarum* and *Hfx. volcanii*.

**S5 Table.**

Thresholded Cytoscape input files

**S1 File.**

**Supplementary Methods:** Supplementary methods file that details simulation construction and parameter choice.

## Data Availability

Code for GRN-BMuSeR and for all downstream analysis is available through Github (https://github.com/andrew-soborowski/GRN BMuSeR). RNA-seq data generated for this project are publicly available via NCBI GEO at accession number GSE338384.

## Acknowledgments

This work was also supported by grants from the National Science Foundation (NSF MCB 2427099 and NSF 1936024) and the National Institutes of Health (NIH 1R35GM158161-01) to A.K.S. The funders had no role in the study design, data collection and analysis, decision to publish, or preparation of the manuscript.

The authors thank Richard Bonneau and Bonneau lab members for initial conversations in getting the project started. We are indebted to the technical support of Sierra Deleon for cell culturing, RNA extractions, and sample organization for RNA-seq data for this project. We thank the Duke Sequencing and Genomic Technologies core facility for their technical support with sequencing the RNA.

