## Supplementary Methods for "A Bayesian Multi-Species Approach Infers Gene Regulatory Networks Across Non-Model Organisms"

### Full_Supplemental_Methods

#### Supplemental Methods: Simulations

Network simulations were constructed as follows. For each simulation, an ancestral network consisting of 100 genes and 300 interactions was initialized (see Ancestral Network Construction). 20 genes were selected to be transcription factors by allowing them to have out-edges, while the remaining 80 genes were constrained to only having in-edges. 10% of TFs (2) were selected as high-impact TFs, corresponding to master regulators, while the remaining 90% (18) were selected to be low-impact TFs, corresponding to TFs that regulate a small subset of genes of specific function. High-impact TFs were parameterized to have an average of 5 times as many out-edges as low-impact TFs. This ratio was selected to balance an increase in out-degree among certain TFs with the constraint of available target genes and the edge budget. Edge counts for each TF were predetermined, with high-impact TFs assigned the average number of interactions for their class and low-impact TFs having counts sampled from a Poisson distribution given the remaining edge budget. A small number of edges were randomly added or removed as needed to ensure that the edge budget was exactly met.

Ancestral edges were assigned in a two-step process. First, each gene was connected to an in-edge from one random low-impact TF to ensure that every gene was regulated by at least 1 TF. The probability that a given low-impact TF is chosen is proportional to the remaining number of edges the TF has left to assign. Next, for each TF we assign the remainder of its out-edges randomly across the other genes, disallowing self-regulation.

To generate two species networks from the ancestral network, we simulate evolution of each network governed by a target percent divergence (see Subroutine Evolve). We split the ancestral network into two identical copies and applied the following to each network independently. We randomly remove a percentage equal to half-divergence of TFs and non-TFs from each network and remove any edges connected to the removed nodes to simulate gene loss (see Prune Network). We also removed a percentage equal to half divergence of the remaining interactions in each network to simulate the loss of interactions due to mutations (see Prune Edges). We then created new genes and TFs in each network to replace the lost ones, simulating gene gain and duplication events (see Regenerate Edges). Finally, we allowed new interactions to form up to the edge budget. These interactions are biased toward forming between either a new gene or TF, to ensure they are fully connected, but are also allowed to form new connections between old genes and TFs, simulating network rewiring. We identify genes and TFs as homologs between the two networks only if neither was pruned during the evolution step, and we identify edges as homologous only if both the TF and the gene are homologous.

Edge weights, corresponding to the relative strength of the interaction, were generated as independent draws from an inverse gamma distribution with shape parameter = 0.5 and scale parameter = 0.25 (see Generate Edge Weights). Non-homologous edges were randomly assigned as activating or repressing. Homologous edges were randomly assigned as both activating or both repressing in each network with probability equal to half of the divergence parameter.

Expression datasets were simulated from each network as follows (see Generate Dataset). For each condition, expression of each TF was sampled from a standard normal distribution. If the condition is a knockout condition, a random, previously unselected TF was set to expression of 0, while if the condition is an environmental condition, TF expression levels remain unchanged. For simplicity, TFs were not modeled as targets in this simulation. For each target gene, expression was calculated by summing across each TF-target edge multiplied by the corresponding TF expression multiplied by a probability that the interaction is observable in this condition. The probability an interaction is observed models the assumption that TF expression is not always linked to gene expression in each condition, due to factors such as post-translational modifications and environmental sensing. For the results presented here, we set this parameter to 0.4, the number of environmental conditions in each species to 40, and the number of independent trials at each divergence level to 20.

### Pseudocode

Ancestral Network Construction

1. Graph G=(V,E) for vertices V={1..100} and edges E={}
2. Let the first 20 vertices be labeled as TFs, TFs={1..20}, and vertices {1,2} be labeled as high impact TFs (HITFs), HITFs = {3..20}. Let all other TF vertices be labeled as low impacts TFs (LITFs), LITFs=3..20}
3. For each HITF, assign out-edges such that each HITF receives 5 times as many as the average LITF $\frac{300}{2+\frac{18}{5}} = 53$
4. For each remaining LITF, calculate the ratio of remaining edges to LITFs ($e_{LITF}$}) and assign an out-edges to each LITF according to $edges\ Poisson(e_{LITF})$
5. If $\sum assigned\_edges > 300$, randomly remove edges until $\sum assigned\_edges = 300$, such that each TF retains at least 1 out-edge
6. If $\sum assigned\_edges < 300$, randomly add edges until $\sum assigned\_edges = 300$
7. For gene v in V, assign an edge from an LITF, $litf$. Choose the LITF with probability proportional to the unassigned edges associated with each LITF and such that $g \neq litf$
8. For each $tf$ in TFs, randomly assign edges from $tf$ to genes in V, such that each edge is unique and such that there is no edge $e(tf, tf)$

Subroutine Evolve

1. Given ancestral network G and divergence parameter d
2. Let A, B, and H be duplicates of E
3. Prune Network (V, A, H, d)
4. Prune Network (V, B, H, d)
5. prune edges (A,d)
6. regenerate edges (A,d)

Prune Network (V, A, H, d)

1. Randomly select $80 * \left(\frac{d}{200}\right)$ non-TF vertices from $\{21..100\} \subset V$ and $20 * \left(\frac{d}{200}\right)$ TF vertices from $\{1..20\} \subset V$
2. For each selected vertex v, remove any edge e from A such that for any edge e(a,b), a$\neq$v and $b \neq$v
3. For each selected vertex v, remove any edge e from H such that for any edge e(a,b), a$\neq$v and $b \neq$v

Prune Edges (A, H, d)

1. For each edge $e(a, b) \in A$, remove $e(a, b)$ from A with probability = $\frac{d}{200}$
2. If $e(a, b)$ was removed from A and $e(a, b)$ exists in H, remove $e(a, b)$ from H

Regenerate Edges (V, A)

1. For each gene v in V, if there is no edge $e(a, b)$ in A such that $b = v$, create edge $e(a, v)$ where $a \in V$ is a LITF and $a \neq v$
2. For each $tf$ in TFs in V, if there is no edge $e(a, b)$ in A such that $a = tf$, create edge $e(tf, b)$ where $b \in V$ and $tf \neq b$
3. Calculate remaining unassigned edge budget, $E_u$, as $E_u = 300 - |E|$
4. Calculate HITF out-edges to assign as $E_{HITF} = 53 * |HITFs| - |e = (a, b) \in E|a \in HITFs|$ and $LITF$ out-edges to assign as $E_{LITF} = E_u - E_{HITF}$
5. For each edge $e(a, b)$ in $E_{HITF}$, randomly choose $a \in HITF$ with probability of selecting a given a inversely proportional to the square of the number of out edges from a. Then, choose b randomly such that b$\neq$a.
6. For each edge $e(a, b)$ in $E_{LITF}$, randomly choose $a \in LITF$ with probability of selecting a given $a$ inversely proportional to the square of the number of out edges from $a$. Then, choose $b$ randomly such that $b \neq a$.

Generate Edge Weights

1. For edge $e(a, b)$ in $A$
    1. Sample edge weight sign $w_{sa}$ uniformly from {-1, 1}

2. Sample edge weight magnitude $w_{ma}$ as a draw from $InvGamma(0.5, 0.25)$
3. Assign edge weight $w_a = w_{sa} * w_{ma}$
4. If edge $e(a, b)$ in H and B
    1. Calculate second edge magnitude $w_{sb}$ as $w_{sa}$ * x∈ {-1,1} with $p\left(\frac{d}{200}, 1 - \frac{d}{200}\right)$
    2. Sample second edge weight magnitude $w_{mb}$ as a draw from $InvGamma(0.5, 0.25)$
    3. Assign edge weight $w_b = w_{sb} * w_{mb}$
5. For edge e(a,b) in B
    1. If edge e(a,b) has no calculated weight
        1. Sample edge weight sign $w_{sb}$ uniformly from {-1, 1}
        2. Sample edge weight magnitude $w_{mb}$ as a draw from $InvGamma(0.5, 0.25)$
        3. Assign edge weight $w_b = w_{sb} * w_{mb}$

Generate Dataset

1. For each condition, for each species A,B
    1. For each $tf$ vertex in V, sample $tf$ weight $w_{tf}$ as $N(0, 1)$
    2. For each non-tf vertex in V (v∈ {21,100})
        1. Vertex expression $v_e = \sum_e (w_a * w_{tf} * p_i) + noise$ where $p_i \in 0, 1$ with $p(0.6, .0, 4)$ and $noise \sim N(0, .5)$
