## Supplementary material for "A Bayesian Multi-Species Approach Infers Gene Regulatory Networks Across Non-Model Organisms": Figure S1

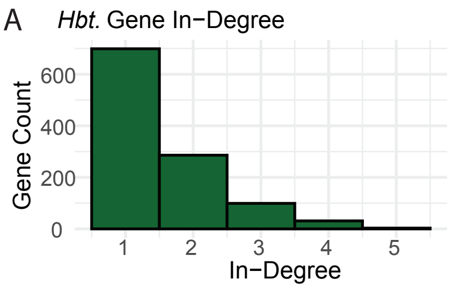


mean1.53
88\% of targets have 2 or fewer TFs as regulators

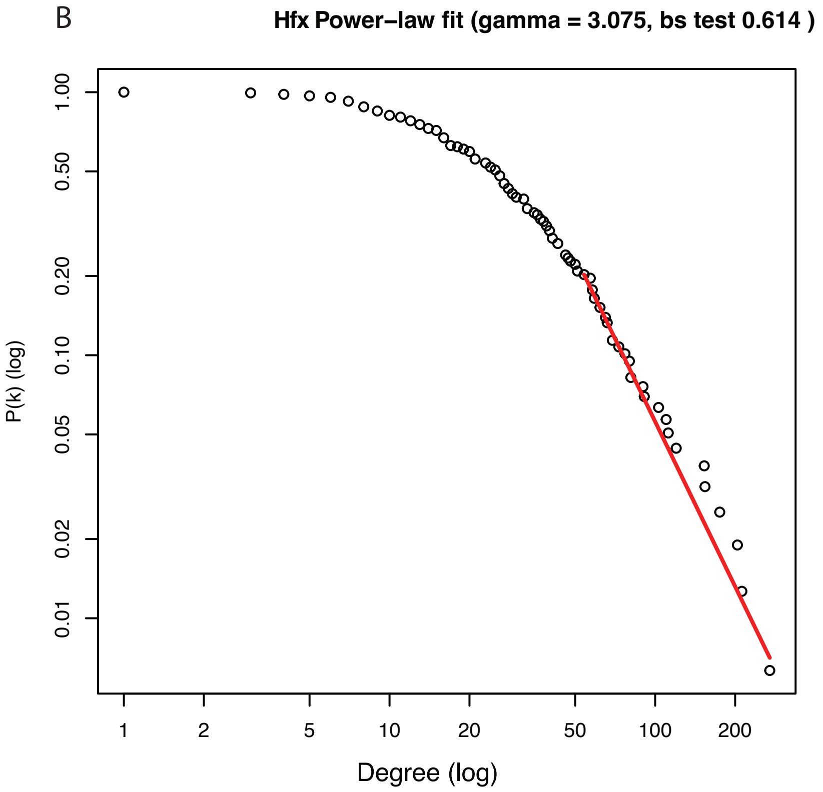


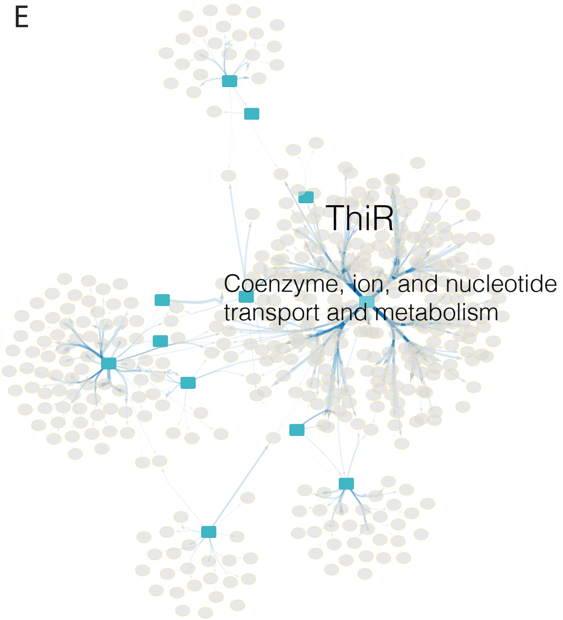


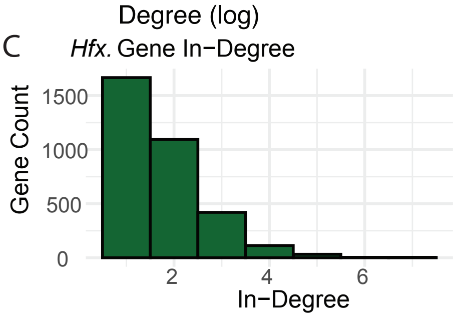


The average in-degree for target genes is 1.73,
with 83\% of targets predicted to have 2 or fewer regulators

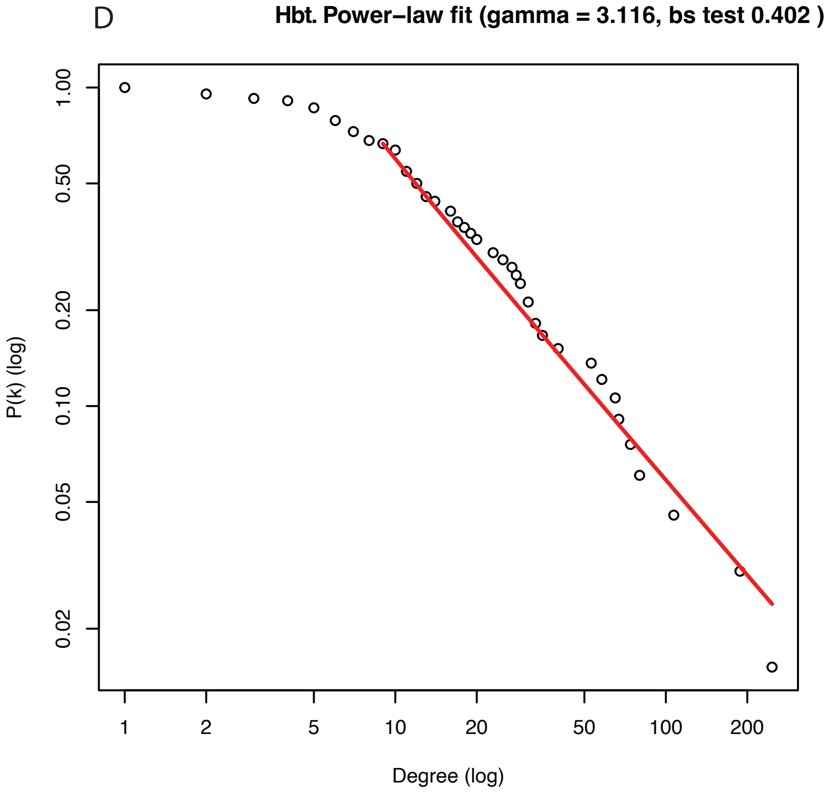
